# Oxidized LDL regulates efferocytosis through the CD36-PKM2-mtROS pathway

**DOI:** 10.1101/2023.09.07.556574

**Authors:** Jue Zhang, Jackie Chang, Vaya Chen, Mirza Ahmar Beg, Wenxin Huang, Lance Vick, Yaxin Wang, Heng Zhang, Erin Yttre, Ankan Gupta, Mark Castleberry, Ziyu Zhang, Wen Dai, Shan Song, Jieqing Zhu, Moua Yang, Ashley Kaye Brown, Zhen Xu, Yan-Qing Ma, Brian C. Smith, Jacek Zielonka, James G. Traylor, Cyrine Ben Dhaou, A Wayne Orr, Weiguo Cui, Ze Zheng, Yiliang Chen

**Author notes:** To whom correspondence should be addressed: Yiliang Chen, Ph.D.; Jue Zhang, M.D., Ph.D.

## Abstract

Macrophage efferocytosis, the process by which phagocytes engulf and remove apoptotic cells (ACs), plays a critical role in maintaining tissue homeostasis. Efficient efferocytosis prevents secondary necrosis, mitigates chronic inflammation, and impedes atherosclerosis progression. However, the regulatory mechanisms of efferocytosis under atherogenic conditions remain poorly understood. We previously demonstrated that oxidized LDL (oxLDL), an atherogenic lipoprotein, induces mitochondrial reactive oxygen species (mtROS) in macrophages via CD36. In this study, we demonstrate that macrophage mtROS facilitate continual efferocytosis through a positive feedback mechanism. However, oxLDL disrupts continual efferocytosis by dysregulating the internalization of ACs. This disruption is mediated by an overproduction of mtROS. Mechanistically, oxLDL/CD36 signaling promotes the translocation of cytosolic PKM2 to mitochondria, facilitated by the chaperone GRP75. Mitochondrial PKM2 then binds to Complex III of the electron transport chain, inducing mtROS production. This study elucidates a novel regulatory mechanism of efferocytosis in atherosclerosis, providing potential therapeutic targets for intervention.

**SUMMARY:** Macrophages clear apoptotic cells through a process called efferocytosis, which involves mitochondrial ROS. However, the atherogenic oxidized LDL overstimulates mitochondrial ROS via the CD36-PKM2 pathway, disrupting continual efferocytosis. This finding elucidates a novel molecular mechanism that explains defects in efferocytosis, driving atherosclerosis progression.

## INTRODUCTION

Professional phagocytes, particularly macrophages, engage in efferocytosis, a crucial physiological process that removes apoptotic cells (ACs) (Arandjelovic and Ravichandran, 2015). This mechanism not only prevents the accumulation of non-functioning cells but also triggers anti-inflammatory and pro-resolving responses in macrophages (Schilperoort et al., 2023b). Defects in efferocytosis often lead to secondary necrosis of ACs, promoting unresolved chronic inflammation and driving the progression of atherosclerosis (Yurdagul et al., 2017). In an atherogenic environment, multiple factors contribute to increased cellular apoptosis, including oxidative stress, hyperlipidemia, and mitochondrial dysfunction (Li et al., 2022). This necessitates more rapid and continual efferocytosis by the surrounding macrophages, especially where ACs significantly outnumber macrophages. While significant progress has been made on the mechanisms of efferocytosis during atherosclerosis (Yurdagul et al., 2017), the regulatory mechanisms of continual efferocytosis remain incompletely understood.

During the initiation stage of atherosclerosis, apoB-containing lipoproteins retained in the arterial subendothelial space are modified into atherogenic particles, including oxidized LDL (oxLDL) (Moore and Tabas, 2011). These particles subsequently bind to macrophage surface receptors such as CD36, promoting intracellular lipid accumulation and foam cell formation (Chen et al., 2015; Rahaman et al., 2006). We previously reported that oxLDL/CD36 signaling reprograms macrophage metabolism from mitochondrial respiration to glycolysis, accompanied by increased fission and production of mitochondrial reactive oxygen species (mtROS) (Chen et al., 2019). Using a diet-induced atherosclerosis *Apoe*-null mouse model (Nakashima et al., 1994), we further showed that mtROS levels are continuously elevated in circulating monocytes in a CD36-dependent manner during the initiation of atherosclerosis. Suppressing mtROS using a transgenic strategy (<u>m</u>itochondrial-specific human <u>cat</u>alase overexpression, or MCAT) protected mice from early atherogenesis (Wang et al., 2014), indicating a contributing role of mtROS. While we and others have revealed connections between mtROS and pro-inflammatory signaling, (Chen et al., 2019; Wang et al., 2014), two important questions remain: 1. Do mtROS modulate macrophage cellular functions such as efferocytosis? and 2. What is the mechanism of mtROS induction by oxLDL?

In this study, we first tested the hypothesis that mtROS are involved in a positive feedback mechanism for continual efferocytosis (where primary efferocytosis promotes secondary efferocytosis). We then investigated the pathological role of oxLDL in the dysregulation of efferocytosis via mtROS. To uncover the underlying molecular mechanisms of mtROS induction, we conducted a combination of transcriptomics, proteomics, immunological, and biochemical assays. Our findings elucidate a novel mechanism underlying impaired efferocytosis during atherosclerosis progression, providing potential therapeutic targets for intervention

## RESULTS

### Macrophage mtROS Facilitate Continual Efferocytosis

When murine peritoneal macrophages were incubated with ACs, a subpopulation of macrophages displayed elevated mtROS production (Figure 1A), as measured by mtROS-specific indicator, MitoNeoD (Shchepinova et al., 2017). The induction of mtROS in response to ACs prompted us to investigate whether mtROS play a role in sustaining continual efferocytosis. We designed a two-stage experimental protocol to investigate the continual efferocytosis capability of macrophages (Figure 1B). In the primary (1^st^) efferocytosis assay, macrophages were co-incubated with PKH67-labeled apoptotic cells (ACs) for 15 minutes. In the secondary (2^nd^) efferocytosis assay, macrophages were first co-incubated with non-labeled ACs for 1 hour. Afterward, unbound ACs were washed out. Macrophages were then co-incubated with PKH67-labeled ACs for 15 minutes. After each stage, we performed extensive PBS washes to remove unbound labeled ACs. Flow cytometry was then used to quantify PKH67 signals from the macrophages. The comparison between primary and secondary efferocytosis served as an indicator of the macrophages’ capacity for continual efferocytosis. Meanwhile, we compared mitochondrial-catalase overexpressed macrophages (MCAT) (Wang et al., 2014) with WT macrophages (Figure 1B). In WT macrophages, secondary efferocytosis was 40% higher than primary efferocytosis. MCAT macrophages displayed similar primary efferocytosis capacity to WT macrophages. However, the enhancement of secondary efferocytosis was fully abolished, suggesting that mtROS are important in the positive feedback mechanism that promotes continual efferocytosis (Figure 1C). The difference in continual efferocytosis is unlikely due to reduced efferocytosis receptors such as MerTK (Thorp et al., 2008), as MCAT macrophages exhibited a 39% higher surface expression of MerTK (Figure S1A).

**Figure 1.**
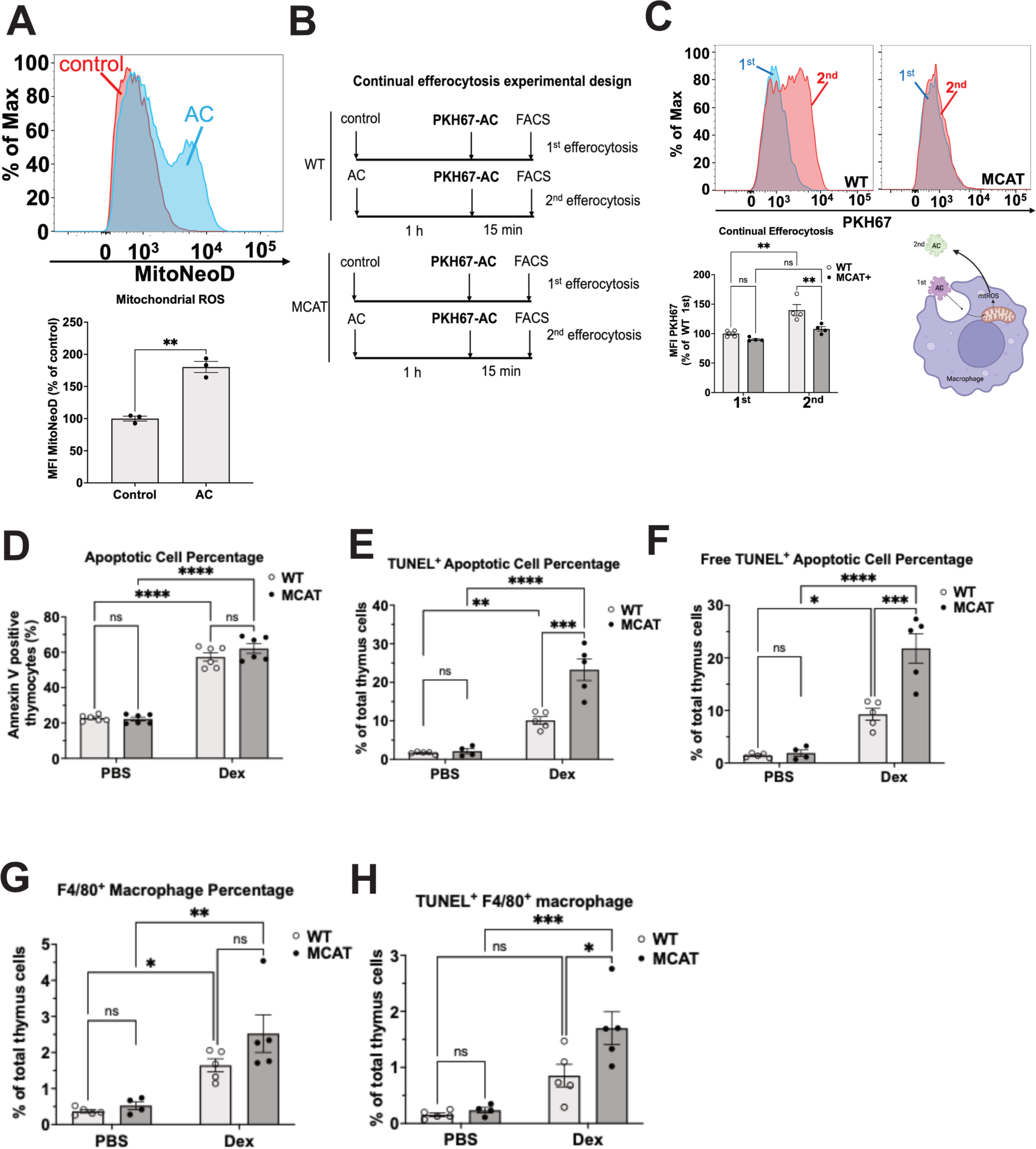
Macrophage mtROS Facilitate Continual Efferocytosis. **A**, Examples of histograms of MitoNeoD fluorescence in WT peritoneal macrophages co-incubated with AC for 30 min, followed by MitoNeoD staining. The MitoNeoD mean fluorescence intensity (MFI) was quantified and shown in the bar graph below; n=3 per group. **B**, The schematic experimental design for the continual efferocytosis assay. **C**, Examples of histograms of PKH67 fluorescence in WT and MCAT macrophages. The PKH67 MFI was quantified and shown in the bar graph below; n=4 per group. The proposed model of how mtROS provide a positive feedback mechanism for continual efferocytosis is shown on the bottom right. **D**, WT, or MCAT thymocytes were treated with 5 μg/ml dexamethasone (Dex) for 20 h *in vitro*, followed by PE-conjugated Annexin V staining for PS-exposed AC. AC rates among all thymocytes were quantified by flow cytometry and shown in the bar graph; n=6 per group. **E-H**, WT, or MCAT mice were peritoneally injected with dexamethasone (250 μg in 1 ml PBS per mouse). 20 h later, mice were sacrificed, and thymocytes were stained with TUNEL and anti-F4/80 antibody, followed by flow cytometry analysis. The percentages of total TUNEL^+^ cells (**E**), free TUNEL^+^ cells (TUNEL^+^/F4/80^-^) (**F**), total F4/80^+^ macrophages (**G**), and TUNEL^+^/F4/80^+^ efferocytes (**H**) were quantified and shown in the bar graph. Max, maximum fluorescence intensity. ns, not significant; *, p<0.05; **, p<0.01; ***, p<0.001; ****, p<0.0001.

To investigate the role of mtROS in modulating efferocytosis for tissue homeostasis, we employed an established *in vivo* efferocytosis assay in mouse thymi (Park et al., 2011; Wang et al., 2017). WT and their littermates MCAT mice were injected intraperitoneally with dexamethasone to induce thymocyte apoptosis. Annexin V binding assays confirmed significant increases in apoptotic cell rates in both groups. No difference was observed between WT and MCAT thymocytes (Figure 1D). 20 hours post-injection, mouse thymi were harvested and TUNEL staining was performed to label ACs together with F4/80 immunostaining (macrophage marker) prior to flow cytometry analysis. The gating strategies for single-cell selection and the dot plots from the double staining are shown in Figure S1B. In PBS (vehicle control)-injected mouse thymi, TUNEL^+^ AC rates are very low for both WT (1.7%) and MCAT (2.1%) groups (Figure 1E), consistent with the notion that efferocytosis is highly efficient *in vivo* and AC are quickly cleared by surrounding macrophages (Yurdagul et al., 2017). Interestingly, while dexamethasone increased TUNEL^+^ AC rates in both groups, AC rates doubled in MCAT thymi (23.3%) compared to those in WT thymi (10.1%) (Figure 1E). A similar pattern was observed for the free TUNEL^+^ AC (defined as TUNEL positive but F4/80 negative) rates (Figure 1F), which indicated that suppressed mtROS induction reduced AC clearance. The difference in AC clearance was unlikely due to different numbers of thymic macrophages, as F4/80^+^ thymic macrophages were comparable between WT and MCAT mice (Figure 1G). Moreover, it was also unlikely due to less recognition of AC in MCAT thymic macrophages because even more efferocytes (defined as TUNEL and F4/80 double positive cells) were observed in dexamethasone-injected MCAT thymuses (Figure 1H). Taken together, our results indicated that macrophage mtROS facilitated efferocytosis *in vivo*, possibly through a positive feedback mechanism.

### OxLDL Impairs Continual Efferocytosis through CD36 and mtROS

We previously reported that oxidized LDL (oxLDL) highly stimulated mtROS through CD36 in macrophages (Chen et al., 2019). To investigate the impact of oxLDL on efferocytosis, we pre-treated WT murine macrophages with 20 μg/ml of lipoproteins for 24 hours, followed by a primary efferocytosis assay. Flow cytometry showed that oxLDL, but not LDL or HDL, increased 3.7-fold PKH67 signals in WT macrophages (Figure 2A). This effect was dose-(Figure S2A) and time-dependent (Figure S2B). To test whether this effect was dependent on types of ACs, we used murine neutrophils isolated from whole blood and allowed them to undergo spontaneous apoptosis before labeling with PKH67 for the primary efferocytosis assay. Consistently, more PKH67 signals were detected in oxLDL-pretreated macrophages (Figure S2C). Furthermore, oxLDL had a similar impact on primary efferocytosis in human monocyte-derived macrophages (HMDMs) (Figure S2D). Taken together, the atherogenic oxLDL elevated primary efferocytosis in murine and human macrophages regardless of the types of ACs.

**Figure 2.**
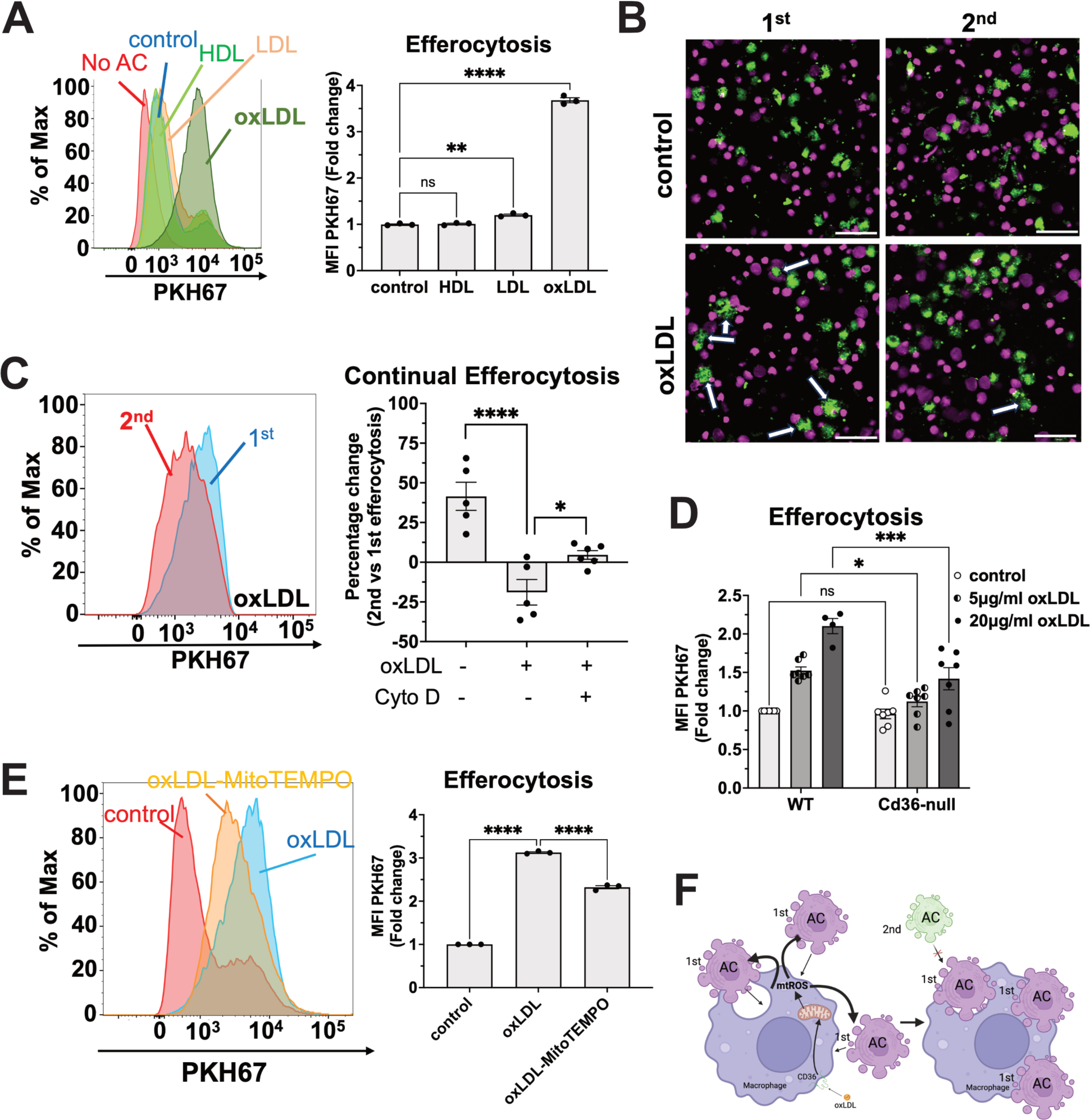
OxLDL Stimulates Efferocytosis through CD36 and mtROS. **A**, Examples of histograms of PKH67 fluorescence in WT peritoneal macrophages pre-treated with 20 μg/ml HDL or LDL or oxLDL for 24 h, followed by PKH67-labeled AC co-incubation for 15 min and flow cytometry analysis. The PKH67 MFI was quantified and shown in the bar graph; n=3 per group. **B**, Representative live-cell confocal images from WT peritoneal macrophages with or without 20 μg/ml oxLDL 24 h pre-treatment. Then, cells were subjected to continual efferocytosis assay (1^st^ vs. 2^nd^), and images were taken at the end of the assay. Efferocytes were labeled in green and ACs were labeled in purple. Scale bar: 50 μm. White arrows point to efferocytes simultaneously engaging more than one AC. **C**, WT peritoneal macrophages were pre-treated with 20 μg/ml oxLDL for 24 h before continual efferocytosis assay in the presence or absence of 5 μM cytochalasin D. Examples of histograms of PKH67 fluorescence in 1^st^ and 2^nd^ efferocytosis with oxLDL pretreatment are shown. The PKH67 MFI was quantified and shown in the bar graph; n=5-6 per group. **D**, WT, or *Cd36*-null peritoneal macrophages were pre-treated with 5 or 20 μg/ml oxLDL for 24h before co-incubation with PKH67-labeled AC for 15 min as primary efferocytosis assay. The PKH67 MFI was quantified and shown in the bar graph; n=3-4 per group. **E**, Examples of histograms of PKH67 fluorescence in WT peritoneal macrophages pre-treated with 20 μg/ml oxLDL or in combination with 1mM MitoTEMPO for 24 h before primary efferocytosis assay. The PKH67 MFI was quantified and shown in the bar graph; n=3 per group. **F**, The proposed model that oxLDL impairs continual efferocytosis. Max, maximum fluorescence intensity. ns, not significant; *, p<0.05; **, p<0.01; ***, p<0.001; ****, p<0.0001.

To complement our flow cytometry data, we captured confocal images at the end of both the primary and secondary efferocytosis protocols (Figure S2E). Our observations revealed distinct differences between oxLDL-pretreated macrophages and control macrophages. Control macrophages typically contacted one AC at a time during both primary and secondary efferocytosis (Figure 2B and Video S1-S2). However, oxLDL-pretreated macrophages frequently made physical contact with multiple ACs simultaneously during primary efferocytosis (Video S3), but not secondary efferocytosis (Video S4). These findings provide a potential explanation for the observed >3-fold increase in PKH67 signals in oxLDL-pretreated macrophages after primary efferocytosis (Figure 2A). To further elucidate the macrophage behavior during efferocytosis, we next employed a label-free live-cell imaging technique (NanoLive Inc), which allowed us to visualize the cells in real-time. Interestingly, while control macrophages appeared to uptake ACs one by one sequentially (Video S5), oxLDL-pretreated macrophages extended their filopodia and attempted to uptake multiple ACs simultaneously (Video S6), in agreement with our confocal imaging observations (Figure 2B). Despite their increased engagement, oxLDL-pretreated macrophages often failed to fully engulf ACs due to the large size of the ACs (Video S6). This incomplete engulfment may potentially impair continual efferocytosis. To corroborate, we repeated continual efferocytosis assay (Figure 1B) before (control) or after macrophages were pre-treated with oxLDL. While secondary efferocytosis was 41% higher than primary efferocytosis in control macrophages, a 24% reduction in secondary efferocytosis was observed in oxLDL-pretreated macrophages (Figure 2C). Moreover, using an actin polymerization inhibitor, cytochalasin D to block AC internalization, we showed that the reduction in secondary efferocytosis observed in oxLDL-pretreated macrophages was fully eliminated (Figure 2C). These results indicate that oxLDL pretreatment specifically impairs the macrophages’ ability to internalize ACs during continual efferocytosis.

Compared to WT macrophages, *Cd36*-null macrophages displayed comparable PKH67 signals at the basal level (control condition) but showed less increase in response to oxLDL (Figure 2D). Moreover, the mtROS scavenger MitoTEMPO attenuated PKH67 signals induced by oxLDL-pretreatment (Figure 2E). Based on these findings, we propose a model where oxLDL/CD36 signaling triggers mtROS overproduction, leading to altered macrophage behavior. This results in simultaneous engagement with multiple ACs, which paradoxically impairs the efficiency of continual efferocytosis (Figure 2F).

### OxLDL Stimulates Phagocytosis through CD36-mtROS pathway

The behavior of oxLDL-pretreated macrophages suggested an attempt to enhance phagocytosis. To test this hypothesis, we conducted *in vitro* phagocytosis assays using pHrodo-conjugated *E. coli* bioparticles. The pHrodo is non-fluorescent outside the cell at neutral pH but brightly fluorescent in acidic phagolysosomes. We quantified pHrodo fluorescence intensity using flow cytometry as an indicator of phagocytosis efficiency (Neaga et al., 2013). Consistent with efferocytosis results, oxLDL, but not LDL or HDL, increased pHrodo signals within macrophages by 87%, which was fully blocked by the phagocytosis inhibitor cytochalasin D (Figure 3A). OxLDL-stimulated phagocytosis was dose-(Figure S3A) and time-dependent (Figure S3B). Similar stimulation was observed in human monocyte-derived macrophages (HMDMs) (Figure 3B). The oxLDL effect was not limited to the phagocytosis of *E. coli* bioparticles because oxLDL also stimulated the phagocytosis of polystyrene beads (Figure S3C). Since polystyrene beads lack bacteria surface antigens such as LPS, this result suggested that the oxLDL phagocytosis-stimulating effect was not mediated by LPS/TLR4 signaling. As a validation, the phagocytosis-stimulating effect was also observed in *Tlr4*-null macrophages, even more pronounced compared to WT macrophages (Figure S3D). Instead, oxLDL-induced phagocytosis was attenuated in *Cd36*-null macrophages (Figure 3C), indicating that this effect involves CD36. Since oxLDL still increased phagocytosis, although to a lesser extent, in *Cd36*-null macrophages, this effect was unlikely mediated through direct interaction between macrophage surface CD36 and bioparticles. Thus, we reasoned that intracellular signaling cascades downstream of the oxLDL/CD36 axis are likely involved in the oxLDL-stimulated phagocytosis.

**Figure 3.**
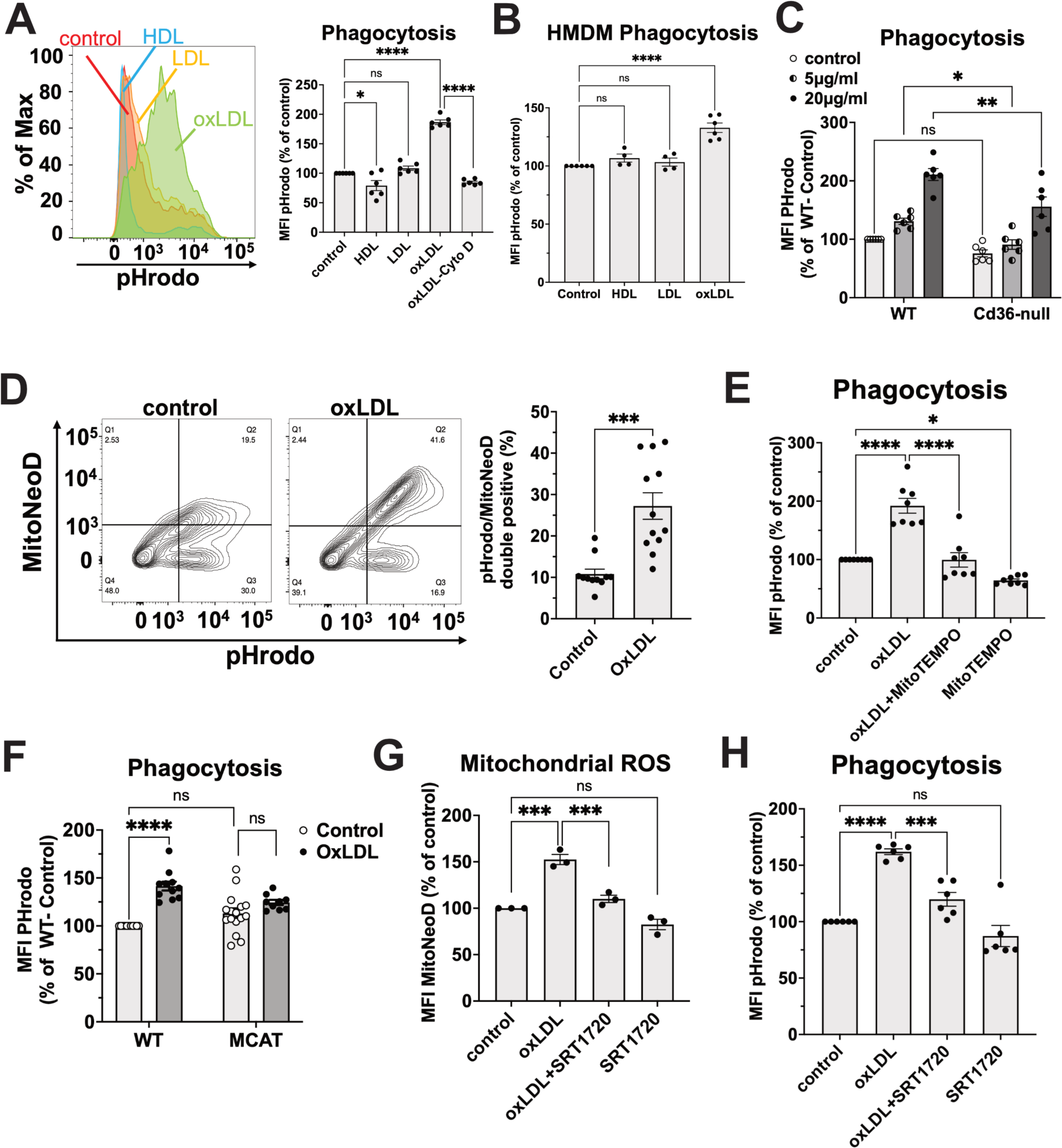
OxLDL Stimulates Phagocytosis through CD36 and mtROS. **A,** Examples of histograms of pHrodo fluorescence in WT peritoneal macrophages pre-treated with 20 μg/ml HDL, LDL, or oxLDL, or oxLDL plus 5 μM cytochalasin D for 24 h, followed by pHrodo-conjugated *E. coli* co-incubation for 30 min and flow cytometry analysis. The pHrodo MFI was quantified and shown in the bar graph; n=3 per group. **B,** HMDMs were pre-treated with 50 μg/ml HDL, LDL, or oxLDL for 24 h before phagocytosis assay. The pHrodo MFI was quantified and shown in the bar graph; n=3 per group. **C,** WT or *Cd36*-null peritoneal macrophages were pre-treated with 5 or 20 μg/ml oxLDL for 24 h before phagocytosis assay. The pHrodo MFI was quantified and shown in the bar graph; n=3 per group. **D,** Examples of contour plots showing MitoNeoD and pHrodo signals from WT peritoneal macrophages pre-treated with 20 μg/ml oxLDL for 24 h. The percentage of MitoNeoD/pHrodo double positive populations was quantified and shown in the bar graph; n=3-5 per group. **E,** WT peritoneal macrophages pre-treated with 20 μg/ml oxLDL for 24 h, followed by 1 mM MitoTEMPO incubation for 1 h before phagocytosis assay. The pHrodo MFI was quantified and shown in the bar graph; n=3 per group. F, WT, or MCAT peritoneal macrophages pre-treated with 20 μg/ml oxLDL for 5 h before phagocytosis assay. The pHrodo MFI was quantified and shown in the bar graph; n=4-5 per group. **G,** WT peritoneal macrophages pre-treated with 20 μg/ml oxLDL for 24 h, followed by 20 μM SRT1720 for 1 h before MitoNeoD staining. The MitoNeoD MFI was quantified and shown in the bar graph; n=3 per group. **H,** The same treatment as in G before phagocytosis assay. The pHrodo MFI was quantified and shown in the bar graph; n=3 per group. Max, maximum fluorescence intensity. ns, not significant; *, p<0.05; **, p<0.01; ***, p<0.001; ****, p<0.0001.

### OxLDL-Induced Phagocytosis Is Dependent on Mitochondrial Reactive Oxygen Species (mtROS)

To investigate the role of mtROS in oxLDL-induced phagocytosis, we co-stained macrophages with MitoNeoD, a specific mitochondria-targeted fluorescent redox probe (Chen et al., 2019; Shchepinova et al., 2017), and pHrodo-conjugated *E. coli* bioparticles. OxLDL pre-treatment significantly increased a macrophage subpopulation with high levels of both mtROS and pHrodo signals (Figure 3D), suggesting a positive correlation between mtROS levels and phagocytosis efficiency. Moreover, MitoTEMPO fully blocked oxLDL-induced phagocytosis (Figure 3E). Consistently, MCAT macrophages were resistant to oxLDL stimulation in phagocytosis, although the basal phagocytosis efficiency was not affected (Figure 3F). Cellular redox status is regulated by a protein lysine deacetylase Sirt1, the activation of which suppresses mtROS production in macrophages (Alam et al., 2021; Singh et al., 2018). We used SRT1720, an allosteric activator of Sirt1 (Mitchell et al., 2014), and showed that SRT1720 significantly reduced both mtROS (Figure 3G) and phagocytosis (Figure 3H) as induced by oxLDL. Taken together, mtROS play a crucial role in mediating oxLDL-induced enhancement of phagocytosis in macrophages.

Using a mitochondrial-targeted redox cycler Mito-nitropyridine (MitoNP), we showed that, although MitoNP promoted mtROS in macrophages (Figure S3E), it inhibited phagocytosis (Figure S3F). Thus, mtROS alone are insufficient to increase phagocytosis efficiency in macrophages.

### Aortic Phagocytic Macrophages Display MtROS Induction during Atherosclerosis

OxLDL promotes foam cell formation and facilitates the initiation and development of atherosclerosis (Chen et al., 2015; Park et al., 2009; Rahaman et al., 2006). We reported that mtROS were continuously elevated in circulating monocytes during the initiation stages of atherogenesis in *Apoe*-null mice (Chen et al., 2019). Circulating monocytes infiltrate the lesion sites and give rise to plaque macrophages (Swirski et al., 2007; Tacke et al., 2007). Thus, we next investigated the mtROS-phagocytosis axis in the aortic lesional macrophages. For *ex vivo* experiments, *Apoe*-null or *Apoe/Cd36* double-null mice were fed a chow diet or HFD for 6 weeks. The aorta were dissected, digested to single-cell suspension, and stained with anti-F4/80 (macrophage marker), anti-Trem2 (aortic foam cell marker) (Cochain et al., 2018), MitoNeoD, and *E. coli*-pHrodo (Figure 4A). Clumped cells were excluded for analysis using the FSC-A vs. FSC-H gating strategy (Figure S4A). As expected, HFD led to increased aortic F4/80^+^ macrophages (Figure S4B) and an 81% increase in F4/80^+^/Trem2^+^ aortic foamy macrophages in *Apoe*-null mice (Figure 4B). Interestingly, the amounts of F4/80^+^/Trem2^+^ aortic foamy macrophages were decreased in *Apoe/Cd36* double-null mice under both chow and HFD conditions (Figure 4B), consistent with a previous report that *Apoe/Cd36* double-null mice were protected from HFD-induced atherosclerosis (Febbraio et al., 2000). 6-week of HFD resulted in a 53% increase in mtROS levels in aortic foamy macrophages from *Apoe*-null mice. However, HFD-induced mtROS were not observed in *Apoe/Cd36* double-null mice (Figure 4C), suggesting that CD36 is required for mtROS induction under atherogenic conditions. Moreover, HFD led to a 25% increase in phagocytosis efficiency in aortic foamy macrophages from *Apoe*-null mice but not in those from *Apoe/Cd36* double-null mice (Figure 4D).

**Figure 4.**
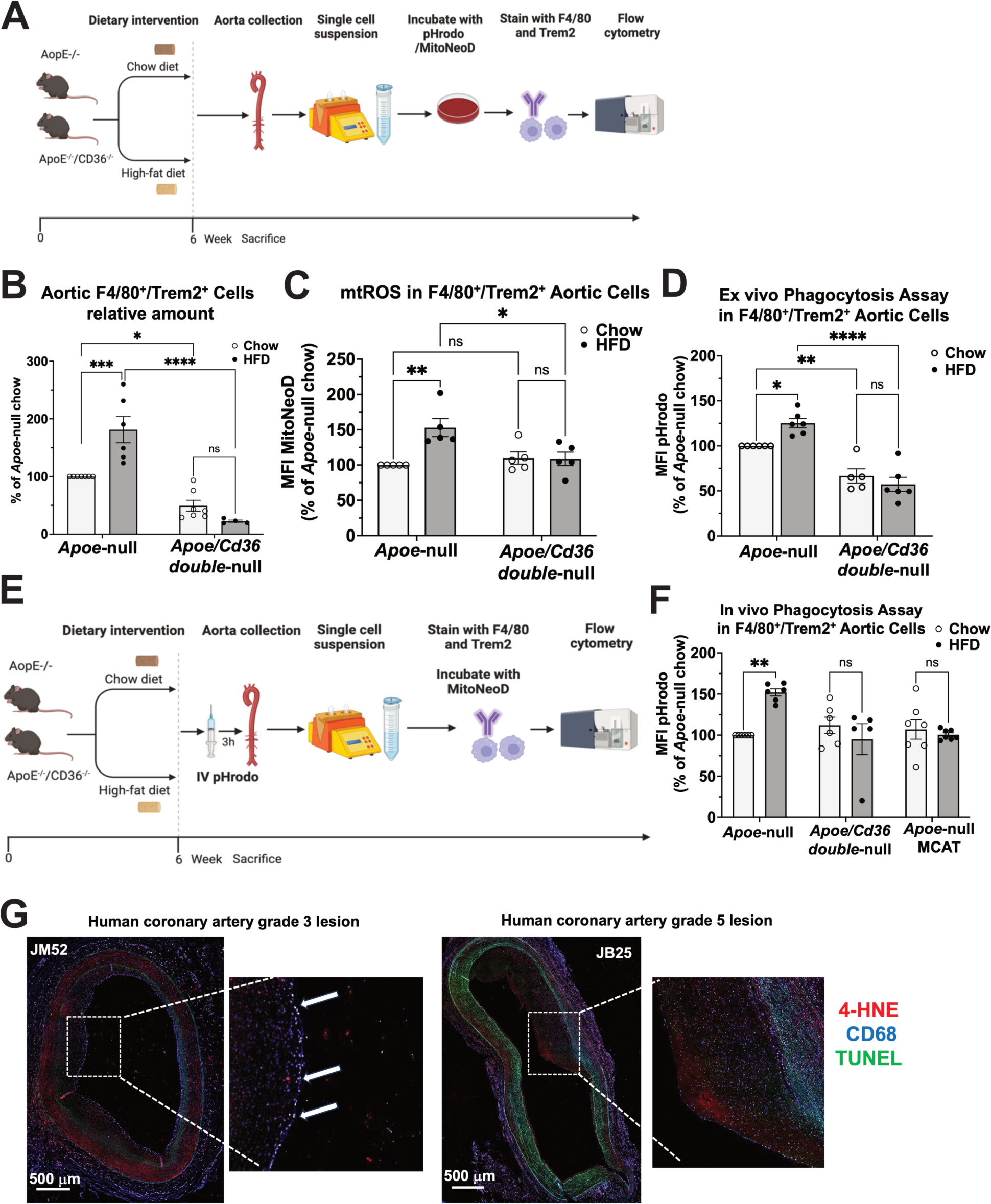
Aortic Phagocytic Macrophages Are Associated with MtROS during Atherosclerosis. **A,** A diagram of the *ex vivo* mtROS and phagocytosis assay. **B,** Total amount of aortic F4/80^+^/Trem2^+^ macrophages were quantified and shown in the bar graph; n=4-7 individual mice per group. **C,** The MitoNeoD MFI was quantified in aortic F4/80^+^/Trem2^+^ macrophages and shown in the bar graph; n=5 individual mice per group. **D,** The pHrodo MFI was quantified in aortic F4/80^+^/Trem2^+^ macrophages and shown in the bar graph; n=5-6 individual mice per group. **E,** A diagram of the *in vivo* phagocytosis assay. **F,** The pHrodo MFI was quantified in aortic F4/80^+^/Trem2^+^ macrophages and shown in the bar graph; n=5-7 individual mice per group. **G,** Fluorescence images of human coronary arteries with grade 3 (left) or grade 5 atherosclerosis lesions triple stained with CD68 (blue), 4-HNE (red), and TUNEL (green). The white rectangle area within the left image was magnified and shown on the right for both samples. White arrows point to the three-color colocalized positions (white). Scale bar: 500 μm. ns, not significant; *, p<0.05; **, p<0.01; ***, p<0.001; ****, p<0.0001.

We consolidated the *ex vivo* results by conducting an *in vivo* phagocytosis assay. We injected *E. coli*-pHrodo through tail veins, waited 3 h, and collected mouse aorta for flow cytometry analysis (Figure 4E). Consistently, HFD increased phagocytosis efficiency by 52% in *Apoe*-null mice, but not in *Apoe/Cd36* double-null mice or *Apoe*-null/MCAT mice (Figure 4F). Moreover, the HFD-stimulated phagocytosis effect was only displayed in F4/80^+^/Trem2^+^ macrophages from the aorta but not in other tissues, including adipose tissue and the liver (Figure S4C). Additionally, peripheral blood monocytes had no detectable pHrodo signal (data not shown). Thus, induction of mtROS contributed to an increase in phagocytosis efficiency in aortic foamy macrophages in a CD36-dependent manner during the initiation of atherosclerosis in mice.

Next, we investigated the association of ROS signaling and efferocytosis in human atherosclerotic plaques. We acquired fixed human atherosclerotic plaque samples (Table S1) and performed three-color co-staining with anti-4-HNE (red), a ROS biomarker (Shoeb et al., 2014), macrophage marker anti-CD68 (blue), and AC marker TUNEL (green). To validate the use of 4-HNE as a marker of oxLDL-induced mtROS signaling, we showed that oxLDL led to high 4-HNE staining in cultured WT peritoneal macrophages but not in MCAT macrophages (Figure S4D). In human grade 3 atherosclerotic lesions, 4-HNE signals co-localized with macrophage and AC markers at the lesion surfaces (shown as white dots and pointed by white arrows in Figure 4G), suggesting a positive association between ROS signaling and efferocytes in early lesions. Interestingly, this surface co-localization was less observed in advanced stage grade 5 lesions (Figure 4G and Figure S4E-S4G). This observation aligns with previous reports of defective efferocytosis in advanced lesions (Stary et al., 1995; Yurdagul et al., 2017). Additionally, as strong colocalization was observed immediately facing the blood vessel lumen, it suggests that ACs may be derived from circulating blood cells, although the exact cell type remains to be identified.

### OxLDL Stimulates Pyruvate Kinase Muscle 2 (PKM2) Mitochondrial Translocation, Which Induces mtROS

We next investigate the molecular mechanism underlying oxLDL-induced mtROS production. We hypothesize that a second messenger downstream of the oxLDL/CD36 complex delivers the signal from the cytoplasm to the mitochondria, inducing mtROS. We previously demonstrated that oxLDL induced a metabolic switch from mitochondrial oxidative phosphorylation to glycolysis (glycolytic switch) (Chen et al., 2019). To test the potential role of glycolytic switch in mtROS induction, we used the glycolysis inhibitor 2-deoxy-D-glucose (2-DG) (Wick et al., 1957) and showed that 2-DG significantly blocked both oxLDL-induced mtROS and phagocytosis efficiency (Figure S5A), suggesting that activation of the glycolytic enzymes may be involved.

If a glycolytic enzyme contributes to mtROS induction under atherogenic conditions, we reason that the expression pattern of that enzyme among different aortic immune cells may be correlated with mtROS levels. Therefore, we re-analyzed published single-cell RNA sequencing (scRNA-seq) data on CD45^+^ immune cells isolated from mouse aorta during atherogenesis (Cochain et al., 2018). Using the same markers for distinct immune cell sup-types, we identified 11 cell clusters shown by uniform manifold approximation and projection (UMAP) (Figure 5A). The aortic macrophage foam cell population was identified as Trem2^high^ cluster based on the previous studies (Cochain et al., 2018; Kim et al., 2018) and shown in the UMAPs (Figure 5A and 5B). Interestingly, the *Pkm* gene, which encodes the last rate-limiting step of the glycolysis, was highly expressed in both Ly6c^high^ pro-inflammatory monocytes and the Trem2^high^ foam cells (Figure 5C and 5D). This was correlated with elevated mtROS levels from the circulating Ly6c^high^ pro-inflammatory monocytes (Chen et al., 2019) and F4/80^+^/Trem2^+^ aortic foamy macrophages (Figure 4C). Nevertheless, genes (i.e., *Hk1*, *Hk2*, H*k3, Pfkm, Pfkl, Pfkp*) that encode the other two rate-limiting steps of glycolysis showed no such correlation (Figure S5B). Moreover, the *Pkm* highly expressed cells appeared predominantly from the HFD-fed mice (Figure 5C and Figure S5C), suggesting upregulation of *pkm* expression in the plaque-forming immune cells during atherogenesis. As a further validation, we reanalyzed another published bulk RNA-seq data on foamy vs nonfoamy macrophages from mice atherosclerotic plaques (Kim et al., 2018). Consistently, *Pkm* expression was significantly elevated in foamy macrophages along with *Cd36* and its known signaling partners (i.e. *Atp1a3* (Chen et al., 2015) and *Itgb3* (Moodley et al., 2003)) and fatty acid trafficking partners (i.e. *Fabp4* and *Cpt1a* (Chen et al., 2019)) (Figure 5E).

**Figure 5.**
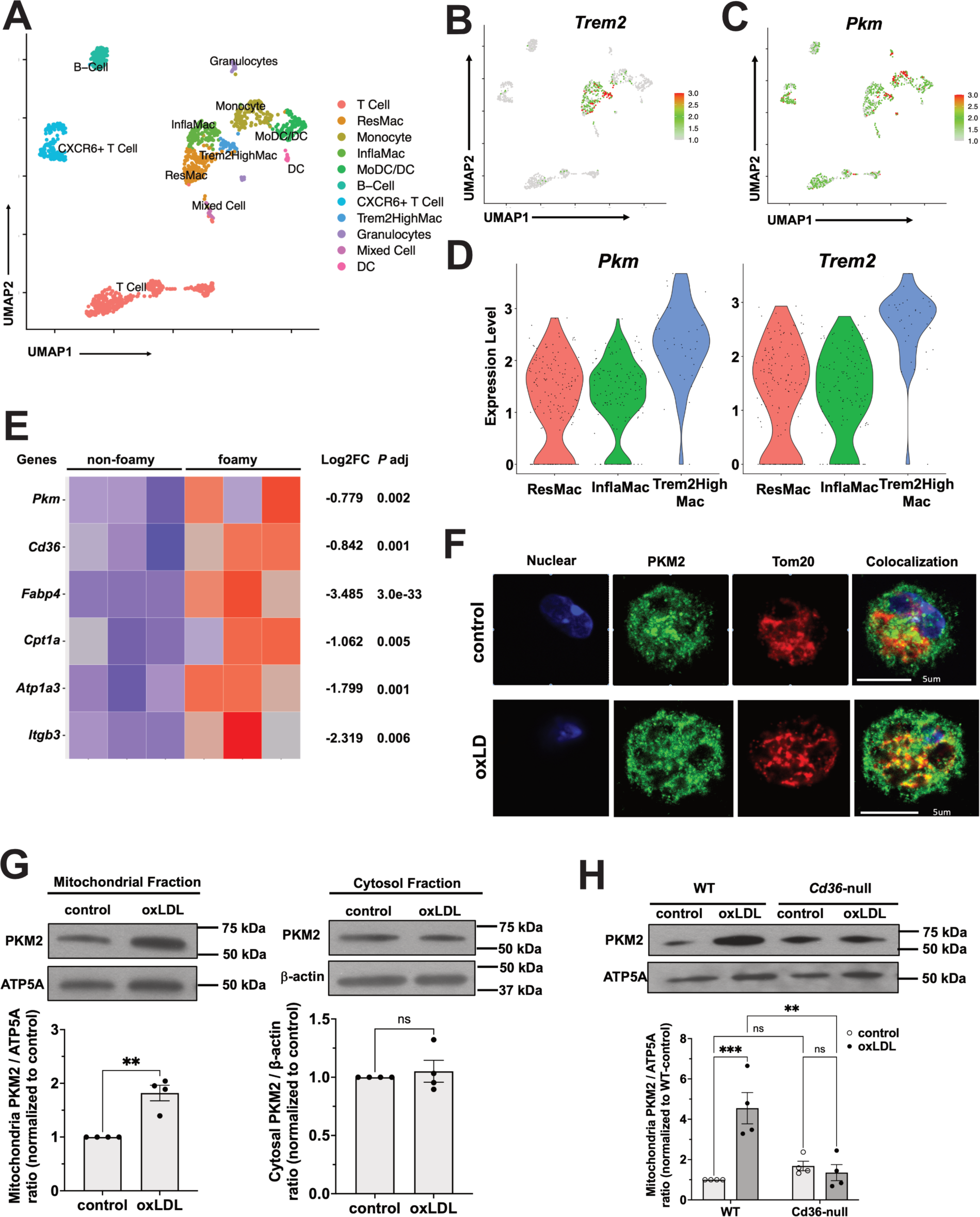
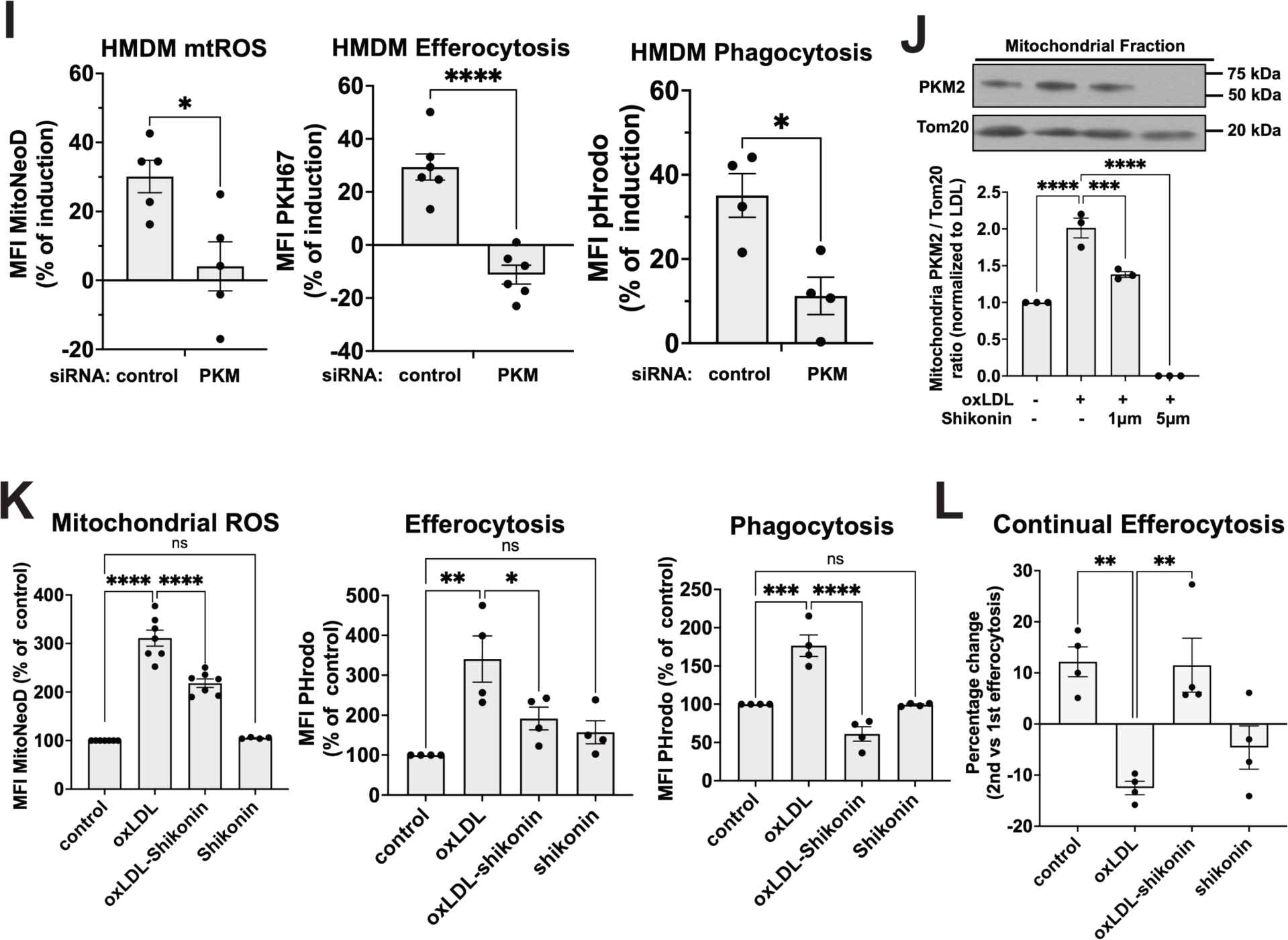
Pyruvate kinase M2 (PKM2) translocate to the mitochondria, promoting mtROS and phagocytosis in macrophages. **A-D**, Mouse scRNA-seq data were re-analyzed from a previous publication (Cochain et al., 2018). Uniform manifold approximation and projection (UMAP) representation of 11 aortic CD45^+^ immune cell clusters are shown in **A**. *Trem2* gene expression pattern (**B**) and *Pkm* gene expression pattern (**C**) are shown in the UMAP. **D**, Violin plots show the *Pkm* and *Trem2* expression distribution among aortic macrophage subpopulations. ResMac: resident macrophages; InflaMac: inflammatory macrophages. **E**, Bulk RNA-seq data were re-analyzed from another publication (Kim et al., 2018). A heatmap is shown of differential expression of relevant genes comparing between foamy and nonfoamy macrophages from mice aorta. **F**, Representative confocal images of macrophages immunostained for PKM2 (green) and Tom20 (red). Nuclei were stained by DAPI (blue). Scale bar: 5 μm. **G**, HMDMs treated with 20 μg/ml LDL (control) or oxLDL for 3 h were lysed and subjected to cell fractionation into mitochondrial and cytosol fractions. PKM2 and ATP5A (mitochondria fraction loading control) blot images from mitochondrial fractions were shown on the left. PKM2 and β-actin (cytosol fraction loading control) blot images from cytosol fractions are shown on the right. Images were quantified, normalized to each loading control, and expressed as fold change of control. n=4 per group. **H**, WT or *Cd36*-null peritoneal macrophages treated with 20 μg/ml LDL (control) or oxLDL for 3 h and then processed as in F. Mitochondrial fractions were immunoblotted for PKM2 and ATP5A and blot images are shown. Images were quantified, normalized, and expressed as fold change of control. n=4 per group. **I**, HMDMs transfected with control or PKM siRNA were treated with 50 μg/ml oxLDL for 24 h before being subjected to mtROS assay (left panel), primary efferocytosis assay (middle panel), or phagocytosis assay (right panel). MFI was quantified and shown as the percent of induction by oxLDL in the bar graph; n=4-6 per group. **J**, WT macrophages treated with 20 μg/ml oxLDL or pre-treated 1 or 5 μM shikonin before addition of oxLDL, incubating for 3 h, and then processed as in F. Mitochondrial fractions were immunoblotted for PKM2 and Tom20 and blot images are shown. Images were quantified and expressed as fold change of control. n=3 per group. **K**, WT macrophages pre-treated with 20 μg/ml oxLDL or in combination with 0.5 μM shikonin for 24 h before mtROS assay (left panel) or efferocytosis assay (middle panel) or phagocytosis assay (right panel). MFI was quantified and shown in the bar graph; n=3-4 per group. **L**, The same treatment as in K, macrophages were subjected to continual efferocytosis assay. Values are shown as percentage change comparing between 2^nd^ efferocytosis and 1^st^ efferocytosis. ns, not significant; *, p<0.05; **, p<0.01; ***, p<0.001; ****, p<0.0001.

The gene PKM encodes two different isoforms, pyruvate kinase muscle 1/2 (PKM1/2), due to RNA splicing. Since oxLDL upregulated PKM2 but not PKM1 (Figure S5D), we focused on PKM2 in our following mechanistic studies. Although PKM2 is known as a cytosolic enzyme, it was reported that oxLDL stimulated its nuclei translocation (Kumar et al., 2020). Interestingly, besides promoting PKM2 nuclei localization, oxLDL increased PKM2 co-localization with the mitochondria marker Tom20 (Figure 5F). To validate, we employed a mitochondrial/cytosol fractionation method. The purity of each fraction was confirmed by immunoblot on markers of cytosol, mitochondria, and nuclei (Figure S5E). Consistently, oxLDL nearly doubled mitochondrial PKM2 levels in HMDMs, while no change in cytosolic PKM2 levels was observed (Figure 5G). Time course experiments showed that mitochondrial PKM2 levels doubled 1 hour after oxLDL treatment, peaked at 3 hours, and stayed high for up to 24 hours (Figure S5F). However, oxLDL-stimulated PKM2 mitochondrial translocation was not observed in the *Cd36*-null murine macrophages (Figure 5H), indicating that PKM2 is downstream of oxLDL/CD36 signaling. To test the role of PKM2 in mtROS induction, we used siRNA to knock down PKM2 levels in HMDMs (Figure S5G). The knockdown of PKM2 significantly attenuated oxLDL-induced mtROS, primary efferocytosis and phagocytosis (Figure 5I). To further consolidate our hypothesis that PKM2 mitochondrial translocation is responsible for oxLDL-induced mtROS, we used the PKM2 targeting inhibitor shikonin (Chen et al., 2011) and showed that shikonin reduced oxLDL-induced PKM2 mitochondrial translocation (Figure 5J), but not nuclei translocation (Figure S5H). In addition, shikonin significantly inhibited oxLDL-induced mtROS, primary efferocytosis, and phagocytosis (Figure 5K). Furthermore, shikonin rescued continual efferocytosis efficiency in oxLDL-treated macrophages (Figure 5L).

### GRP75 Mediates PKM2 Translocation to the Mitochondria, Where It Binds to the Electron Transport Chain (ETC) Complex III

To elucidate the molecular mechanisms of PKM2 mitochondrial translocation and identify potential targets, we conducted a mass spectrometry analysis of PKM2-interacting proteins. WT peritoneal macrophages were treated with 20 μg/ml oxLDL for 3 hours, lysed, and PKM2 was immunoprecipitated (IP) by a monoclonal rabbit IgG against PKM2. The immunoprecipitates from control and oxLDL-treated cell lysates (n=3) were separated by SDS-PAGE, and gel slices were analyzed by mass spectrometry. An irrelevant monoclonal rabbit IgG IP sample and a blank SDS-PAGE gel served as negative controls. After excluding proteins present in the negative controls, we identified 26 mitochondrial proteins that co-IP with PKM2 (Table S2).

Among the identified proteins, GRP75, a member of the HSP70 family, stood out. GRP75 functions as a chaperone, shuttling between the cytoplasm and mitochondria, and facilitates the import of cytosolic proteins into the mitochondrial matrix (Craig, 2018; D’Silva et al., 2004). Notably, the levels of GRP75 in PKM2 immunoprecipitates from oxLDL-treated samples were 3-fold higher than those from control samples (Table. S2), suggesting an enhanced PKM2/GRP75 interaction in response to oxLDL. We corroborated with PKM2/GRP75 co-IP experiments in both murine macrophages and HMDMs. OxLDL significantly enhanced the PKM2/GRP75 interaction, peaking at 2-3 hours post-treatment (Figure 6A). A control IgG antibody did not precipitate either PKM2 or GRP75, confirming the specificity of the interaction (Figure S6A). Furthermore, immunostaining revealed increased colocalization of PKM2 and GRP75 following oxLDL treatment (Figure 6B). To assess the importance of GRP75’s chaperone function in PKM2 mitochondrial translocation, we employed a GRP75 selective inhibitor MKT-077 (Wadhwa et al., 2000; Wu et al., 2020). Consistently, MKT-077 blocked oxLDL-induced PKM2 mitochondrial translocation (Figure 6C) but showed no effect on nuclei PKM2 (Figure S5H). To validate, HSPA9 (the gene encoding GRP75) siRNA knocked down 65% of GRP75 (Figure S6B) and fully blocked oxLDL-induced PKM2 mitochondrial translocation (Figure 6D). Taken together, the chaperone GRP75 facilitates oxLDL-stimulated PKM2 mitochondrial translocation.

**Figure 6.**
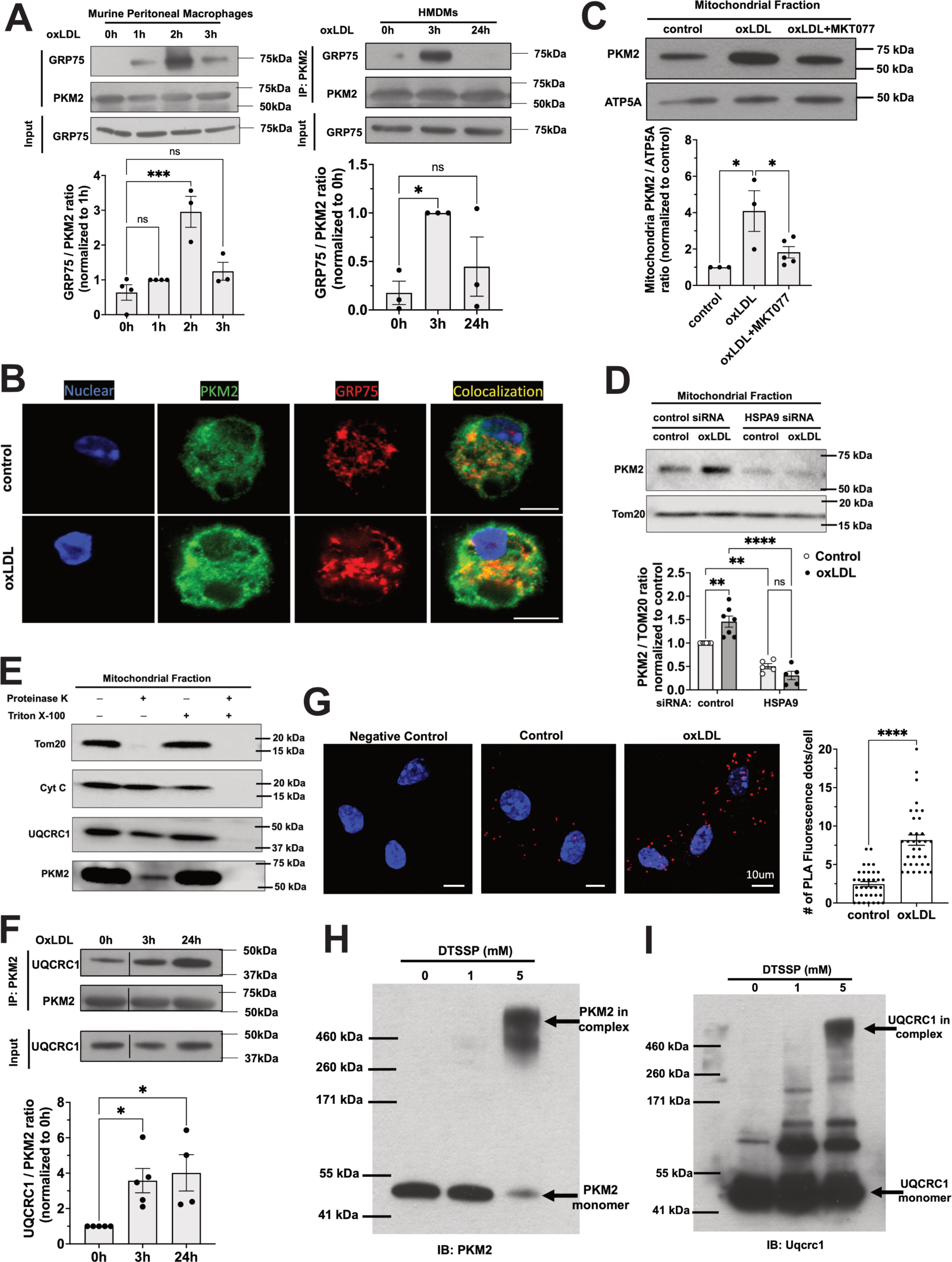
GRP75 mediates PKM2 translocation to the mitochondria, where it interacts with the ETC Complex III. **A**, Murine peritoneal macrophages (left panel) or HMDMs (right panel) were treated with 20 μg/ml oxLDL for indicated time periods before cell lysates were IP with anti-PKM2. Representative Western blot images of GRP75 and PKM2 from PKM2 precipitates and GRP75 from total cell lysates (Input) are shown. Images were quantified and expressed as a fold change of 1 h. n=3 per group. **B**, Representative confocal images of macrophages immunostained for PKM2 (green) and GRP75 (red). Nuclei were stained by DAPI (blue). Scale bar: 5 μm. **C**, WT macrophages were treated with 20 μg/ml oxLDL or pre-treated with 0.1 μM MKT077 before addition of oxLDL, incubating for 3 h before cell fractionation. Mitochondrial fractions were immunoblotted for PKM2 and ATP5A and blot images are shown. Images were quantified and expressed as fold change of control. n=3-5 per group. **D**, HMDMs transfected with control or HSPA9 siRNA were treated with 50 μg/ml oxLDL for 24 h before being the mitochondrial fractions were subjected to immunoblot of PKM2 and Tom20. Images were quantified and expressed as fold change of control. n=5-7 per group. **E**, OxLDL-pretreated WT macrophage mitochondria fractions were subjected to proteinase K digestion in the absence or presence of Triton X-100. Mitochondrial lysates were then subjected to immunoblot of Tom20, cytochrome C, UQCRC1, and PKM2. Representative blot images from four separate experiments were shown. **F**, WT peritoneal macrophages treated with 20 μg/ml oxLDL for indicated time periods before cell lysates were IP with anti-PKM2. Representative Western blot images of UQCRC1 and PKM2 from PKM2 precipitates and UQCRC1 from total cell lysates (Input) are shown. Images were quantified and expressed as fold change of 0 h (control). n=4-5 per group. **G**, Representative confocal images from *in-situ* proximity ligation assay (PLA) show enhanced PKM2/UQCRC1 interaction with or without 20 μg/ml oxLDL 24 h treatment. The PLA signals (red fluorescence dots) were quantified and shown in the bar graph. n=3. The PLA assay without primary antibody addition serves as a negative control. Scale bar: 10 μm. **H**, OxLDL-treated macrophage lysates were incubated with DTSSP crosslinker for 1 h at room temperature, followed by SDS-PAGE and immunoblot for PKM2. A representative blot image is shown. n=5. **I**, The same treatment as in E, followed by immunoblot for UQCRC1. A representative blot image is shown. n=5. ns, not significant; *, p<0.05; ***, p<0.001; ****, p<0.0001.

To determine whether PKM2 was non-specifically attached to the mitochondrial outer membrane or located within the mitochondria, we conducted a proteinase K digestion assay as previously established (Wegrzyn et al., 2009). Mitochondria isolated from oxLDL-treated macrophages were incubated with proteinase K in the presence or absence of triton X-100. Our findings showed that cytochrome C (intermembrane protein) and UQCRC1 (matrix protein) were resistant to proteinase K digestion, while Tom20 (outer membrane protein) was not protected by the mitochondria membranes and degraded. Importantly, a portion of PKM2 was resistant to digestion, demonstrating its presence inside the mitochondria, protected by the mitochondrial membranes. As a validation, addition of triton X-100 compromised mitochondrial membrane integrity, resulting in degradation of all proteins (Figure 6E). These results confirm that a fraction of PKM2 enters the mitochondria rather than merely attaching to the outer membrane, providing strong evidence for the intramitochondrial localization of PKM2 stimulated by oxLDL.

ETC Complex III is one of the major sites of mtROS production (Mailloux, 2015). Our mass spectrometry analysis identified UQCRC1, a subunit of ETC Complex III, exclusively in oxLDL-treated PKM2 immunoprecipitates (Table S2). To validate this finding, we performed PKM2 immunoprecipitation followed by UQCRC1 immunoblotting. As expected, we detected increased UQCRC1 protein levels in PKM2 immunoprecipitates, starting at 3 hours and peaking at 24 hours after oxLDL treatment (Figure 6F). Moreover, we performed an *in-situ* proximity ligation assay (PLA), which corroborated that oxLDL enhanced PKM2/UQCRC1 interaction (Figure 6G). To confirm the direct interaction between PKM2 and ETC Complex III, we employed a crosslinking approach. OxLDL-treated macrophage lysates were incubated with the crosslinking reagent DTSSP prior to SDS-PAGE and immunoblot analysis. In the absence of DTSSP, only PKM2 monomer was detected. However, treatment with 5 mM DTSSP resulted in a significant reduction of the monomer band and the appearance of a high molecular weight band (Figure 6H). This high molecular weight band is consistent with the size of the ETC Complex III (490 kDa) plus PKM2. To exclude the possibility that the high molecular weight band represented a PKM2 oligomer, we performed a control experiment using purified PKM2. The purified protein was subjected to crosslinking with DTSSP under the same conditions. No high molecular weight band was detected in this case (Figure S6C). As a corroboration, immunoblot for UQCRC1 showed a similar molecular weight band in the presence of 5 mM DTSSP (Figure 6I). Collectively, these results provide strong evidence that mitochondrial PKM2 directly binds to ETC Complex III, suggesting a potential mechanism for mtROS induction.

## DISCUSSION

The oxLDL/CD36 signaling axis has been well-established as a key driver of pro-atherogenic phenotypes in macrophages, including foam cell formation (Chen et al., 2015; Rahaman et al., 2006), oxidative stress (Chen et al., 2019), and chronic inflammatory responses (Stewart et al., 2010). We previously demonstrated that this signaling pathway reprograms macrophage metabolism, promoting aerobic glycolysis and mtROS induction, which contribute to atherosclerosis in mice (Chen et al., 2019). In this study, we elucidate the molecular mechanisms linking glycolysis elevation to mtROS induction. Furthermore, we demonstrate that oxLDL-induced mtROS alter macrophage efferocytosis behavior and ultimately impairs continual efferocytosis. Our findings reveal a novel pathological mechanism underlying defective efferocytosis in atherogenic conditions and identify potential new therapeutic targets for atherosclerosis treatment.

In a normal human body, billions of cells undergo programmed cell death and are cleared through efferocytosis every day (Bianconi et al., 2013). This large number is determined by physiological processes such as immune cell selection during development (Opferman, 2008), cell aging (Antonelou et al., 2010) or cells reaching their lifespan (McCracken and Allen, 2014). Pathological processes like tissue damage and regeneration also contribute to this turnover (Juban and Chazaud, 2021). Despite the vast number of apoptotic cells (ACs) generated, they rarely accumulate due to the rapid clearance by surrounding phagocytes, particularly macrophages (Doran et al., 2020). Calculating the precise ratio of ACs to macrophages in a tissue is challenging due to high variability influenced by macrophage status, the source and trigger of cell death, and the local microenvironment. However, it is estimated that ACs can sometimes outnumber macrophages. This imbalance necessitates continual efferocytosis—the capacity of macrophages to rapidly engulf multiple ACs—to maintain tissue homeostasis (Morioka et al., 2019). In this work, we demonstrate that ACs induce mtROS in macrophage, which in turn facilitate continual efferocytosis, creating a positive feedback loop (Figure 1). This mechanism elucidates the high efficiency of efferocytosis and highlights the involvement of mtROS in essential physiological functions. Our findings align with clinical trials showing increased mortality associated with antioxidant supplementation (Bjelakovic et al., 2007), underscoring the beneficial effects of ROS signaling (Sena and Chandel, 2012). However, our subsequent findings reveal a paradoxical effect: while moderate levels of mtROS facilitate efferocytosis, overstimulation of mtROS under atherogenic conditions impairs this process, potentially contributing to the accumulation of ACs in atherosclerotic lesions.

Atherosclerosis exemplifies a chronic disease with mitochondrial dysfunction and excess ROS production, leading to oxidative stress and widespread cell death (Madamanchi and Runge, 2007). Efficient and continual efferocytosis is crucial for removing ACs before they undergo secondary necrosis, which can trigger chronic inflammation and drive atherosclerosis. However, the presence of necrotic cores in advanced human atherosclerotic plaques suggests impaired efferocytosis as the disease progresses (Kolodgie et al., 2004). Despite extensive research into the regulatory mechanisms of efferocytosis, the specific factors disrupting this process within atherosclerotic lesions remain poorly understood (Yurdagul et al., 2017).

Both oxLDL and its receptor CD36 accumulate within human atherosclerotic lesions (Fukuchi et al., 2002; Nakagawa-Toyama et al., 2001). In the present study, we uncover a novel pathological mechanism whereby the oxLDL-CD36-mtROS axis alters macrophage behavior during continual efferocytosis (Figure 2). Under normal conditions, macrophages engage ACs one at a time, uptaking them sequentially (Video S1, S2, and S5). This sequential approach is logical given that ACs can be similar in size to macrophages, necessitating the complete processing of one AC before engaging another. However, oxLDL-pretreated macrophages exhibit altered behavior, simultaneously engaging multiple ACs. This simultaneous engagement ultimately impairs the efficient engulfment of ACs and disrupts continual efferocytosis (Figure 2 and Video S2, S3, and S6). Since this is an *in vitro* observation, whether it also applies to lesional macrophages *in vivo* is unclear. However, we hypothesize that this simultaneous engagement behavior may represent an adaptive response aimed at enhancing phagocytosis, regardless of the nature of the large particles to be engulfed. This hypothesis is supported by both our *in vitro* and *in vivo* assays (Figures 3 and 4). The observed behavior could be interpreted as a compensatory mechanism triggered by oxLDL exposure, potentially aimed at increasing the overall phagocytic capacity of macrophages. However, in the context of efferocytosis, this non-selective approach appears to be counterproductive. This hypothesis aligns with previous observations that apoptosis is atheroprotective in early stages of atherosclerosis (Gautier et al., 2009), when the ratio of ACs to macrophages is relatively low and the demand for continual efferocytosis is minimal. In these early stages, oxLDL-stimulated phagocytosis may be largely beneficial. However, as atherosclerosis progresses, sustained apoptosis in lesional macrophages in advanced stages leads to a significant increase in AC: macrophage ratio (Martinet et al., 2019). In this advanced stage, the oxLDL-stimulated phagocytosis via simultaneous engagement becomes problematic, impairing the efficiency of continual efferocytosis.

The precise mechanism by which oxLDL-induced mtROS lead to simultaneous engagement behavior in macrophages requires further investigation. Our observations reveal that oxLDL pretreatment results in a highly dynamic and distorted macrophage phenotype (Video S6) compared to control conditions (Video S5). These phenotypic changes suggest altered actin cytoskeleton dynamics beneath the cell membrane, a crucial factor in driving the phagocytosis process (May and Machesky, 2001). Recent research has shown that actin can be reversibly modified by ROS, playing a significant role in regulating actin filament dynamics (Rouyere et al., 2022). Furthermore, oxLDL/CD36 signaling has been reported to alter actin dynamics through ROS generation (Park et al., 2009). Based on these findings, we propose a model wherein the oxLDL/CD36 axis regulates actin rearrangement both temporarily and spatially through fine-tuning of mitochondria fission/fusion events, mtROS induction, and modification of actins or actin-associated motor proteins.

A critical question remains as to what initiates mitochondrial dysfunction and ROS induction. Recent studies have shed light on this issue, revealing that initial efferocytosis promotes aerobic glycolysis, which in turn drives continual efferocytosis (Morioka et al., 2018; Schilperoort et al., 2023a). Our findings provide a molecular mechanism for this process, demonstrating that pyruvate kinase M2 (PKM2), a key glycolytic enzyme, translocates to the mitochondria and induces mtROS production (Figures 5 and 6). Unfortunately, the oxLDL-PKM2-mtROS axis appears to be overstimulated, disrupting continual efferocytosis regulation. The role of PKM2 in atherosclerosis is further supported by clinical and experimental evidence. In human patients with coronary artery disease, PKM2 is associated with increased mtROS production (Shirai et al., 2016). Moreover, animal studies have shown that PKM2 contributes to atherogenesis in mice (Doddapattar et al., 2022). Interestingly, in that study, lesional macrophages from myeloid-specific PKM2 deficient mice displayed higher efferocytosis. Moreover, those animals developed smaller necrotic cores, supporting a role of PKM2 in defective efferocytosis during atherosclerosis. Another recent study has demonstrated that oxLDL upregulates PKM2 and stimulates its translocation to the nuclei, where it activates the HIF-1α pathway and promotes aerobic glycolysis as a positive feedback mechanism (Kumar et al., 2020). Our findings extend this understanding by revealing that PKM2 also translocates to the mitochondria, inducing mtROS production. These mtROS likely further stabilize nuclear HIF-1, reinforcing the metabolic shift. These findings highlight PKM2 as a critical mediator in the cellular response to oxLDL exposure, linking oxidative stress, metabolic reprogramming, and mitochondrial dysfunction in the context of atherosclerosis.

Further investigation is needed to elucidate the precise mechanism by which oxLDL/CD36 signaling induces PKM2 mitochondrial translocation. PKM2 subcellular localization is known to be regulated by its oligomeric state and post-translational modifications (Alves-Filho and Palsson-McDermott, 2016). Therefore, it would be valuable to examine whether CD36 downstream signaling triggers specific post-translational modifications of PKM2 that promote its mitochondrial translocation. Our findings provide experimental support for this hypothesis. We demonstrated that shikonin, a well-characterized PKM2 inhibitor, effectively blocked oxLDL-induced PKM2 mitochondrial translocation, mtROS production, and impaired efferocytosis (Figure 5). Shikonin is known to reduce PKM2 phosphorylation and inhibit its dimerization and tetramerization (Huang et al., 2022; Zhao et al., 2018). By inhibiting PKM2 phosphorylation and oligomerization, shikonin appears to disrupt this pathway, preserving normal macrophage function and efferocytosis capacity in the presence of oxLDL.

The mechanism by which mitochondrial PKM2 induces ROS production requires further investigation. Our findings demonstrate that PKM2 binds to multiple subunits of the electron transport chain (ETC) complexes (Table S2), but the precise consequences of these interactions remain unclear. One potential hypothesis is that PKM2/ETC complex interactions may disrupt electron transfer, leading to electron leakage and subsequent superoxide generation. However, this proposed mechanism requires further experimental validation.

The precise identity of mtROS involved in the regulation of efferocytosis remains an open question. While MitoNeoD has been proposed as a selective probe for mitochondrial superoxide (Shchepinova et al., 2017), we cannot definitively rule out the involvement of other oxidants. The inhibitory effects observed with the mitochondria-targeted catalase (MCAT) transgene strategy suggest a significant role for hydrogen peroxide, which is generated from superoxide.

In summary, our study elucidates a novel regulatory mechanism of efferocytosis in an atherogenic environment. OxLDL-induced mtROS impair continual efferocytosis by macrophages.

## MATERIALS AND METHODS

### Experimental Animals

All mice used in this study were on the C57BL/6 background. WT mice were from Charles River (#C57BL/6NCrl), *Apoe*-null mice (#002052), *Tlr4*-null (#007227), and MCAT transgenic mice (#016197) were purchased from the Jackson Laboratory. *Cd36*-null and *Apoe/Cd36* double-null were generated as previously described (Febbraio et al., 2000). MCAT mice were crossed with Apoe-null mice to generate *Apoe*-null/MCAT+ for the *in vivo* phagocytosis assays. All mice were kept in a 12-hour dark/light cycle and fed standard chow ad libitum unless indicated otherwise. Animals of both sexes were used, but since we have not observed sex differences in macrophage CD36-PKM2 signaling axis, data were analyzed together. For high-fat diet challenge experiments, adult mice between 16 to 20 weeks of age were randomly divided into two groups: one was continued on a standard chow diet and the other was placed on an atherogenic high-fat diet (HFD) for 6 weeks before assays. All procedures involving live animals were approved by the Institutional Animal Care and Use Committee at the Medical College of Wisconsin.

### Human Artery Sample Processing

Human aorta and coronary artery samples were collected by Dr. Wayne Orr’s lab (Louisiana State University), embedded in paraffin, and atherosclerotic plaques were scored by a pathologist based on the Stary Classification System (Stary et al., 1995). The samples were subjected to deparaffinization, antigen retrieval, and stained using TUNEL assay kit (#ab66110; Abcam), followed by immunostaining with anti-4-HNE (#ABIN873270; Antibodies-online.com) and anti-CD68 (#ab955; Abcam). The secondary antibody Alexa Fluor647-conjugated antibody (#ab150075; Abcam) targeted 4-HNE antibody, and Alexa Fluor405-conjugated antibody (#ab175660; Abcam) targeted CD68 antibody. Fluorescence images were captured using a Nikon Ti2-E widefield fluorescence inverted microscope with Nikon NIS Elements 5.42.03 acquisition software. To quantify, arterial intimal area were determined based on hematoxylin and eosin (H&E) staining. Analysis was performed by an individual blinded to the stages of plaque using the Fiji software.

### Isolation and *In Vitro* Culture of Peritoneal Macrophages

Mice were injected with 2 ml 4% thioglycolate (#T9032; Sigma) in sterile saline intraperitoneally and 4 days later were sacrificed with CO_2_. Peritoneal cavities were flushed, and macrophages were then suspended in 10 ml PBS pre-warmed to 37°C, counted, and centrifuged at 250g for 5 minutes. Cells were re-suspended in the culture media and seeded into culture dishes. Cells were cultured in RPMI media (#10-040-CV; Corning) supplemented with 10% FBS (#10437028; Gibco), 100 U/ml penicillin, and 100 μg/ml streptomycin (#15140122; Gibco) at 37°C in a humidified incubator with 5% CO_2_. Cells were maintained in full media for at least 48 hours before treatment. All subsequent treatments were conducted in RPMI media with 10% FBS.

### Isolation and *In Vitro* Culture of Human Monocyte-Derived Macrophages (HMDMs)

Monocytes were isolated from human buffy coats (provided by Versiti Blood Donor Center) by Ficoll density centrifugation and allowed to differentiate over 7 days *in vitro* to macrophages as described (Chen et al., 2015). HMDMs were cultured in X-ViVo 15 hematopoietic media (#04418Q; Lonza) supplemented with 5% human serum (#H3667; Sigma), 100 U/ml penicillin, and 100 μg/ml streptomycin at 37°C in a humidified incubator with 5% CO_2_. All following treatments were conducted in X-ViVo 15 media in the presence of 5% human serum.

### Preparation of oxLDL

Human LDL (#360-10) were purchased from Lee BioSolutions (MO, USA). LDL were diluted and oxidized as described here. Briefly, oxidation of LDL (0.5 mg/ml) by Cu^2+^ was performed by dialysis vs. 5 μM CuSO_4_ in PBS for 6 hours at 37°C. Oxidation was terminated by adding BHT (40 μM) and DTPA (100 μM). Then the solution was subjected to dialysis against PBS with DTPA (100 μM) to remove Cu^2+^. For quality control, LDL oxidation was confirmed by thiobarbituric acid reactive substances (TBARS) assay using a kit (#ab118970; Abcam). The oxLDL stock solution (0.5 mg/ml) was put in a 15 ml tube with argon gas flushed above and parafilm wrapped around the cap before the tube was stored in a 4 °C fridge until usage.

### Flow Cytometry Assays

Each cell suspension was prepared in 200 μl Flow Buffer (PBS with 5% FBS and 0.1% w/v sodium azide). All flow cytometry experiments were performed with a BD LSR II instrument using FACSDiva software. The optimal compensation and gain settings were determined for each experiment based on unstained and single color-stained samples. Live cells were gated based on cell forward and side scatter signals. Doublets were excluded based on FSC-A vs. FSC-H plots. Flowjo software version 10.8.1 (Tree Star, OR) was used to analyze the data.

### *In Vitro* mtROS Measurements

Cells were seeded into 6-well plates at 0.25 × 10^6^ cells/well. After treatment, cells were incubated with 5 μM MitoNeoD for 15min. Cells were washed with PBS twice, lifted by scraping, and suspended for flow cytometry analysis using the phycoerythrin (PE) channel.

### *In Vitro* Efferocytosis Assays

Foam cell-derived apoptotic cells (AC) were shown in human atherosclerotic lesions (Ball et al., 1995; Hegyi et al., 1996) and considered as the most relevant AC model in this work. To generate foam cell-derived apoptotic cell bodies, WT murine peritoneal macrophages were treated with 50 μg/ml oxLDL for 48 hours. Apoptosis was confirmed by PE-conjugated Annexin V binding assay (#A35111; Invitrogen). AC were lifted by scraping and labeled with a green fluorescent dye PKH67 (#MINI67; Sigma). PKH67-labeled AC were then subjected to continual efferocytosis assay protocol shown in Figure 1B. Then, macrophages were lifted by scraping, suspended in the Flow Buffer and subjected to flow cytometry analysis using the FITC channel. In a second AC model, mouse neutrophils were isolated from C57BL/6 bone marrow cells using the neutrophil isolation kit (#130-097-658; Miltenyi Biotec) and were left overnight at room temperature, during which cells underwent automatic apoptosis. Apoptotic neutrophils were then labeled with PKH67 and subjected to efferocytosis assay.

### Live Cell Confocal Imaging for *In Vitro* Continual Efferocytosis Assay

Murine peritoneal macrophages were seeded into 35 mm imaging dishes (ibidi USA), 0.25 × 10^6^/dish. For 1st efferocytosis, there is no AC pre-treatment before adding DRAQ7-labeled (NBP2-81126; Novus) AC. For 2nd efferocytosis, macrophages were pre-treated with non-labeled AC with the ratio of AC:macrophages (4:1) for 1 hour, and non-attaching ACs were washed away before the addition of DRAQ7-labeled AC. Macrophages were stained with PKH67 for 5 minutes, followed by addition of DRAQ7-labeled AC with the ratio of AC:macrophages (4:1), and confocal timelapse imaging starts immediately after that. Imaging was performed using a Nikon Ti2-E platform with CSU-W1 Yokogawa spinning disk confocal scanner unit. Nikon NIS Elements 5.42.03 software was used to acquire. The objective used was a dry Nikon 20× Plan Apochromat 0.75 NA. Samples were imaged using the Nikon LUN-F Laser unit. During the 50-minute efferocytosis period, cells were maintained using the Tokai hit incubation chamber with 5% CO_2_ and a temperature of 41°C. 17 XY positions were imaged at a no delay interval for 45 minutes, resulting in 65 loops. Samples were excited with the 488 nm laser at 15% power with an exposure of 500 ms and the 525/50 nm emission filter was used, the sample was then excited with the 647 nm laser at 26% power with an exposure of 800 ms and the 685/40 nm emission filter was used. Images were acquired with a Hamamatsu ORCA-Fusion BT CMOS camera with a pixel size of 6.5 um, and z-stacked 3-D images were created for active efferocyte quantification. Active efferocytes are defined as DRAQ7 (ACs) signals within PKH67 (macrophages) signals and counted at the end of the 50-minute co-incubation.

### *In Vivo* Efferocytosis Assays

The assays were conducted based on a previously established protocol (Park et al., 2011; Wang et al., 2017). Briefly, WT and their litter mate MCAT mice were intraperitoneally injected with dexamethasone (#D4902; Sigma) (250 μg per mouse dissolved in 1 ml PBS). 20 hours after injection, mice were euthanized, and thymuses were harvested. Then, thymuses were briefly rinsed in PBS and disaggregated by mechanical forces. The cell suspension was filtered by a 40 μm cell strainer to remove the large debris. The thymocytes were subsequently stained with TUNEL assay kit substrate and FITC-conjugated F4/80 antibody (#123107; Biolegend) at RT for 15 minutes in the dark. Cells were then washed once with Flow Buffer and resuspended in 200 μl Flow Buffer for flow cytometry analysis.

### *In Vitro* Phagocytosis Assays

Cells were seeded into 6-well plates at 0.25 × 10^6^ cells/well. To determine the phagocytosis efficiency of macrophages, cells were incubated with 20 μg/ml pHrodo-conjugated (#P35366; Invitrogen) E. coli bioparticles for 30 minutes. Cells were washed with PBS twice to remove free bioparticles, lifted by scraping and suspended for flow cytometry analysis using the fluorescein isothiocyanate (FITC) channel. For an alternative phagocytosis assay, cells were incubated with 10 μm green fluorescence beads (#G0100; Thermo Scientific) diluted in RPMI1640 medium in the incubator for 16h. The cells were washed with PBS twice to remove free beads, lifted by scraping and suspended for flow cytometry analysis using the FITC channel.

### Efferocytosis Assay Using Murine Neutrophil as Apoptotic Cells

Murine neutrophils were isolated from C57BL/6 mice bone marrow cells using the neutrophil isolation kit (Miltenyi Biotec, #130-097-658). Non-neutrophil cells were depleted by incubating bone marrow cells with biotinylated antibodies and anti-biotin microbeads. Purified neutrophils were stored in Eppendorf tubes on an open bench at room temperature overnight, during which period they underwent apoptosis. Cells were then labeled with PKH67 and co-incubated with macrophages for efferocytosis assays.

### *Ex Vivo* Aortic F4/80^+^/Trem2^+^ Macrophage mtROS Assays and Phagocytosis Assays

Mice were sacrificed after 6 weeks of chow diet or HFD (#TD.88137; Harlan Teklad). The whole aortas were removed and digested in gentle MACSTM C Tubes (Miltenyi Biotec) with an enzyme mixture (Liberase, 0.77 mg/ml; Hyaluronidase, 0.3 mg/ml; Deoxyribonuclease, 0.3 mg/ml; BSA, 1mg/ml; CaCl_2_, in PBS) by a gentle MACSTM Octo Dissociator with Heaters (Miltenyi Biotec) at 37°C for 45 minutes. The cell suspension was then filtered by a 40 μm cell strainer to remove the large debris. For mtROS assays, the filtered single-cell suspension was incubated with 5 μM MitoNeoD in RPMI1640 with 10% FBS at 37°C for 15 minutes, followed by immunostaining with Pacific Blue-conjugated F4/80 antibody (#123123; Biolegend) and APC-conjugated Trem2 antibody (#FAB17291A; R&D Systems) in Flow Buffer at room temperature (RT) for 15 minutes in the dark. Cells were then washed once with Flow Buffer and resuspended in 200 μl Flow Buffer for flow cytometry analysis. For phagocytosis assays, the filtered single-cell suspension was incubated with 20 μg/ml pHrodo-conjugated E. coli bioparticles in RPMI1640 with 10% FBS at 37°C for 30 minutes, followed by immunostaining with PE/Cy5-conjugated F4/80 antibody (#123111; Biolegend) and PE-conjugated Trem2 antibody (#FAB17291P; R&D Systems) in Flow Buffer at RT for 15 minutes in the dark. Cells were then washed once with Flow Buffer and resuspended in 200 μl Flow Buffer for flow cytometry analysis.

### *In Vivo* Aortic F4/80^+^/Trem2^+^ Macrophage Phagocytosis Assays

At 6 weeks on chow/HFD, each mouse was injected with 50 μg pHrodo-conjugated E. coli bioparticles through the tail vein. 3 hours later, mice were sacrificed, and the aortas were processed as described in the *ex vivo* phagocytosis assay. The pHrodo bioparticle incubation step was skipped.

### PKM2 Immunoprecipitation and Mass Spectrometry for Proteomics Analysis

WT murine peritoneal macrophages (2 × 10^6^ cells) were treated with 20 μg/ml oxLDL for 3 hours. Cells were lysed, and 1 mg of protein was incubated with 2 μg anti-PKM2 IgG (#4053; Cell Signaling) at 4°C overnight. Then, 25 μl of A/G agarose beads (#20421; Thermo Scientific) were added and incubated at 4°C overnight. The beads were pelleted at 10,000g for 1 minute at room temperature, washed three times with lysis buffer, boiled in loading buffer, and the supernatant was loaded to SDS-PAGE gel for electrophoresis. The gel was stained by Coomassie Protein Stain (Abcam), and the gel lanes were excised, including three from control PKM2 IP samples and three from oxLDL-treated PKM2 IP samples, a blank gel (negative control) and a control IgG IP sample (negative control). The excised gels were subjected to mass spectrometry proteomics analysis by the Harvard Medical School Taplin Mass Spectrometry Facility.

### Mitochondria/Cytosol Fractionation

Macrophages were subjected to mitochondria/cytosol fractionation using a commercial Mitochondria Isolation kit (#89874; Thermo Scientific) following the manufacturer’s instruction.

### Crude Mitochondria Isolation and Proteinase K Digestion for Mitochondrial Protein Localization Assay

The assay was conducted based on an established protocol (Zhou et al., 2023). To isolate crude mitochondria, WT murine peritoneal macrophages (8 × 10^6^ cells) were treated with 20 μg/ml oxLDL for 3 hours. Cells were lysed and centrifuged at 500g for 3 minutes at 4°C. Supernatant was removed, and pellets were resuspended in PBS and kept at 4°C for all following steps. Suspension was centrifuged at 500g for 3 minutes, and supernatant was removed. Pellets were chilled on ice for 2 minutes and passed through a 27-gauge needle in 1 ml homogenate buffer (Tris-HCl 350 mM pH 7.8, NaCl 250 mM, MgCl_2_ 50 mM) for 25 times. Then centrifuge at 1,200g for 3 minutes and transfer the supernatant into a fresh Eppendorf tube. Centrifuge at 15,000g for 2 minutes and remove the supernatant. Resuspend pellet in 500 μl buffer A (Tris-HCl 10 mM pH 7.4, EDTA 1 mM, Sucrose 320 mM) and 15,000g for 2 minutes. The pellet is crude mitochondria. The crude mitochondria were resuspended in TD buffer (Triza base 49.99 mM, NaCl 274.13 mM, KCl 20.12 mM, Na_2_HPO_4_ 13.95 mM) and divided into four samples. Sample 1: no treatment. Sample 2: add proteinase K 10 μg/mg protein. Sample 3: add 20% Triton X-100. Sample 4: add both proteinase K and Triton X-100. Incubate at room temperature for 1 hour. Add 1 μl 100 mM PMSF to all samples. Add loading buffer for SDS-PAGE and immunoblot analysis.

### Protein Crosslinking Assay

WT murine peritoneal macrophages (8 × 10^6^ cells) were treated with 20 μg/ml oxLDL for 3 hours. The mitochondrial fraction was incubated with DTSSP at room temperature for 1 hour before addition of 1.2 μl stop solution (1M Tris, pH 7.5) to 30 μl solution. Add loading buffer for SDS-PAGE and immunoblot analysis.

### Transfection of siRNA

The siRNAs against human HSPA9, which encodes GRP75: (#hs.Ri.HSPA9.13.1; Sense: 5’-GUAUCAGCAUGUGCAAAUCUUGUTT-3’ and anti-Sense: 5’-AAACAAGAUUUGCACAUGCUGAUACUG) and PKM (#hs.Ri.PKM.13; Sense: 5’-GAUUAUCAGCAAAAUCGAGAAUCAT-3’ and anti-Sense: 5’-AUGAUUCUCGAUUUUGCUGAUAAUCUU-3’) were obtained from IDT and mixed with INTERFERin® transfection reagent (#55-127; Polyplus) for 10 minutes at RT temperature before transfection. After 10 days of differentiation with ∼70% confluency, HMDMs were transfected with siRNA duplexes at 50 nM concentration in serum-free medium for 24 hours and then replaced with fresh full medium for another 24 hours. The scrambled siRNA (Qiagen) was used as the negative control.

### Immunoprecipitation and Immunoblot Analysis

Cells were washed with PBS and lysed with ice-cold CelLytic M Cell Lysis Reagent (#C2978; Sigma) containing protease inhibitor cocktail (#11697498001; Roche) and phosphatase inhibitors (#524625; Sigma). For immunoprecipitation assays, the lysates were pre-cleared with A/G agarose beads (Thermo Scientific) at 4°C for 1 hour. The supernatants containing the same amount of protein (∼1 mg) were incubated with 10 μl primary antibody overnight at 4°C; 25 μl of A/G agarose beads was then added and incubated overnight at 4°C. The beads were washed and boiled in a loading buffer, and the supernatant was loaded into SDS-PAGE gel. The signals were detected with chemiluminescent substrate (#34579; Thermo Scientific) and quantified by ImageJ software. For immunoblotting, protein concentrations were determined by the NanoDrop One/OneC spectrophotometer (#ND-ONE-W; Thermo Scientific), and equal amounts of proteins were loaded in each lane.

### Immunofluorescence Assay

Macrophages were seeded on uncoated glass coverslips, fixed in 4% paraformaldehyde for 15 minutes, permeabilized in 0.2% Triton X-100 for 10 minutes, and blocked with 3% BSA for 1 hour. The cells were incubated with primary antibodies overnight at 4°C and then fluorescence-conjugated secondary antibodies for 3 hours at RT in the dark. Cells were then washed and counterstained with 4’,6-diamidino-2-phenylindole (DAPI) Vectashield mounting medium (#H-1200; Vector Laboratories) and imaged by Olympus FV1000-MPE Laser Scanning Confocal and Multi-photon Microscope using 60 x/1.2NA water objectives.

### *In-Situ* Proximity Ligation Assay (PLA)

Macrophages were fixed and incubated with rabbit anti-PKM2 (#4053; Cell Signaling) and mouse anti-UQCRC1 monoclonal antibodies (#459140; Invitrogen). Cells were then washed and incubated with species-specific secondary antibodies (#DUO92002; Sigma) conjugated to unique DNA strands. Negative control slides were only incubated with secondary antibodies. The oligonucleotides and a ligase were added to form a circular template, which was then amplified and detected using complementary fluorescently labeled probes. Red fluorescent dots representing protein-protein interaction were visualized by a laser confocal fluorescence microscope.

### Label-Free Live Cell Imaging and Video Generation

WT murine peritoneal macrophages were seeded (0.4 × 10^6^ cells/well) in a 35 mm high glass bottom μ-dish (IBIDI Inc). After 24 hours, the cells were treated with 20 μg/ml oxLDL or maintained in the cell culture media (control) for 24 hours. After that, AC were added to each well in a final ratio of 4AC:1macrophage, and the control and oxLDL dishes were immediately placed in the CX-A label-free cell imaging system (Nanolive S.A., Switzerland). The images from a 90 μm × 90 μm area were taken every 15 seconds for 45 minutes. After image acquisition, all images from each condition were combined to generate a short video using ImageJ software. Time-lapse live cell images were processed utilizing Fiji image analysis software, and video annotations were produced using the open-source plugin, draw_arrow_in_movies. Trajectories were manually defined for each annotation and overlayed with time-lapse images. Arrow annotations were color-coded to highlight the presumptive macrophages and apoptotic cells, as determined by visual observation of three immune cell microscopists. The resulting annotated time-lapse images were then processed and exported as graphics interchange format files.

### Statistics

Data are presented as means ± standard error of the mean (SEM), and all statistical analysis was done using GraphPad Prism 9 software. D’Agostino-Pearson omnibus normality test was performed to confirm that data are normally distributed, followed by the Student’s t test (two-tailed, unpaired, and equal variance) for comparisons between two groups. For data comparison involving more than two groups, ANOVA was used followed by Dunnett’s or Tukey’s multiple comparisons test. Statistical values, including number of replicates (n), are noted in the figure legends. In all figures, “ns” means not significant. *P<0.05, **P<0.01, ***P<0.001, ****P<0.0001. For in vitro studies n=number of biological repeats while for the in vivo studies, n=number of animals.

### Online supplemental material

**Figure S1** compares the surface efferocytosis receptor MerTK expression between MCAT and WT, and shows the gating strategies for the *in vivo* efferocytosis assay. **Figure S2** shows additional data supporting that oxLDL stimulates efferocytosis *in vitro*. **Figure S3** shows that oxLDL stimulated phagocytosis *in vitro*, but only inhibiting ETC complex or inducing mtROS is insufficient to stimulate phagocytosis. **Figure S4** shows additional data on *in vivo* phagocytosis assay in mice and human aorta immunostaining and quantification. **Figure S5** shows additional data supporting that PKM2 expression is specifically associated with aortic foamy cells, and its mitochondrial translocation was rigorously tested. **Figure S6** shows addition data supporting the specificity of PKM2 co-IP experiments, and the PKM2-ETC complex interaction. **Table S1** shows the basic information of human artery samples. **Table S2** shows the list of 26 mitochondrial proteins co-IP with PKM2. **Video S1** (control) and **S2** (oxLDL pre-treated) show the macrophage phenotypical dynamics during the efferocytosis process (both tracked for 45 minutes).

### Data Availability

The authors declare that all supporting data and methods are available within the article and the Supplemental Material. The scRNA-Seq data for reanalysis were retrieved from Gene Expression Omnibus (GEO) under accession number GSE97310. The bulk RNA-Seq data for reanalysis were retrieved under the accession number GSE116239. Additional supporting data and methods are available from the corresponding authors upon reasonable request.

## ACKNOWLEDGMENTS

We thank Drs. Michael P. Murphy (University of Cambridge) and Richard C. Hartley (University of Glasgow) for providing MitoNeoD for mtROS measurements. We thank Ms. Kathryn Williams, Ms. Savannah Neu, and Mr. Douglas Franklin for their excellent technical support. Additionally, we thank Dr. Dinesh Jaishanker (Northwestern University) for providing the CX-A imaging system, and Ms. Justine Johnson (Nanolive SA) for helping with the live cell imaging experiments. We also thank our colleagues, Dr. Roy Silverstein, Dr. Daisy Sahoo, and Dr. Mary Sorci-Thomas for their helpful discussion and comments on our work. Some graphs included were created with BioRender.com.

## Non-Standard Abbreviations and Acronyms

AC: apoptotic cell bodies
2-DG: 2-deoxy-D-glucose
ETC: electron transport chain
FSC: forward scatter
HFD: high fat diet
HMDM: human monocyte-derived macrophages
IP: immunoprecipitation
MCAT: mitochondrial-targeted human catalase expression
MFI: mean fluorescence intensity
mtROS: mitochondrial reactive oxygen species
oxLDL: oxidized low-density lipoprotein
PKM2: pyruvate kinase muscle 2
PLA: proximity ligation assay
RT: room temperature
scRNA-seq: single-cell RNA sequencing
Trem2: triggering receptor expressed on myeloid cells 2
TUNEL: terminal deoxynucleotidyl transferase dUTP nick end labeling
UMAP: uniform manifold approximation and projection
WT: wild type

