## Supplementary material for "Oxidized LDL regulates efferocytosis through the CD36-PKM2-mtROS pathway": Fig. s1-s6, Table S1,S2

### SUPPLEMENTAL FIGURES

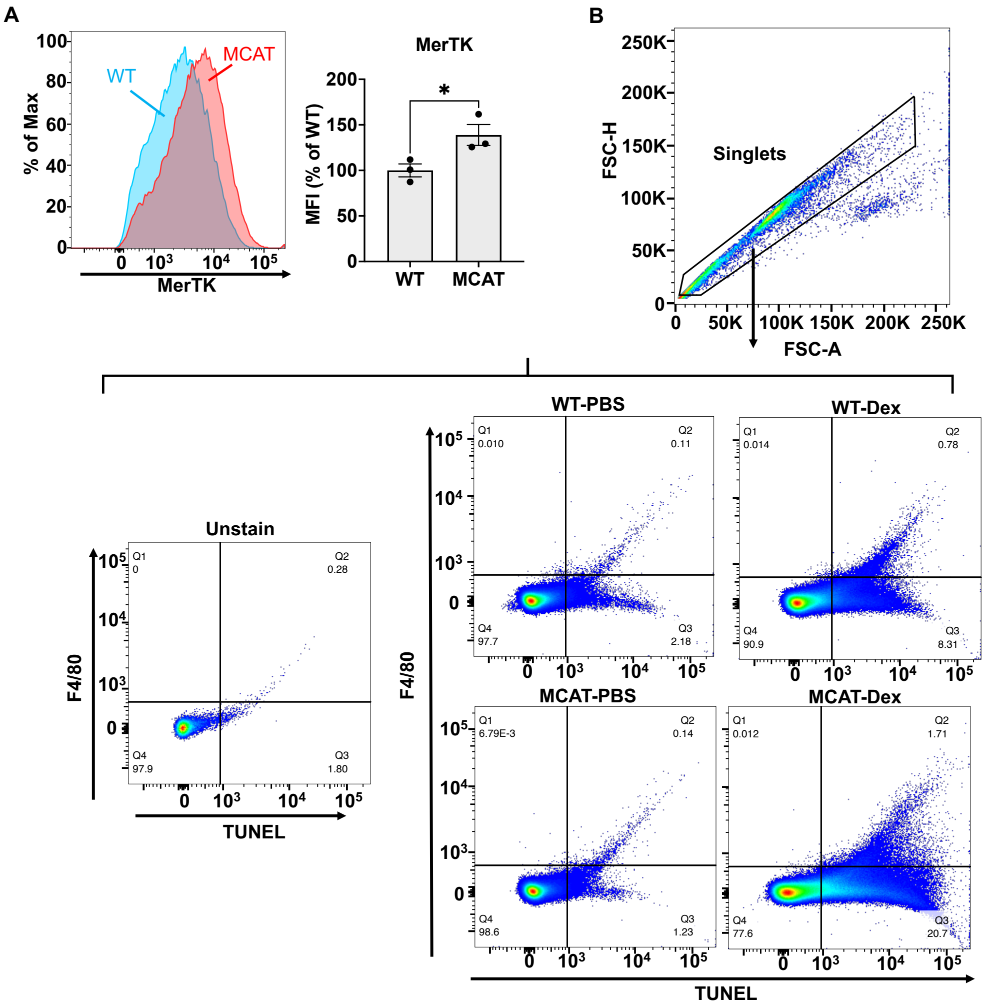


**Figure S1. MCAT transgenic mice show reduced efferocytosis *in vivo*. A,** Examples of histograms of MerTK surface expression in WT and MCAT peritoneal macrophages. MFIs are shown in the bar graph; n=3 per group. **B**, The gating strategies of the *in vivo* efferocytosis assay. Cell clumps were excluded based on FSC-A vs FSC-H dot plots. Following single-cell population selection, examples of dot plots with quadrant images of TUNEL and F4/80 co-staining are shown for all 4 conditions. An unstained sample dot plot is also shown, which was used to determine the quadrant gating.

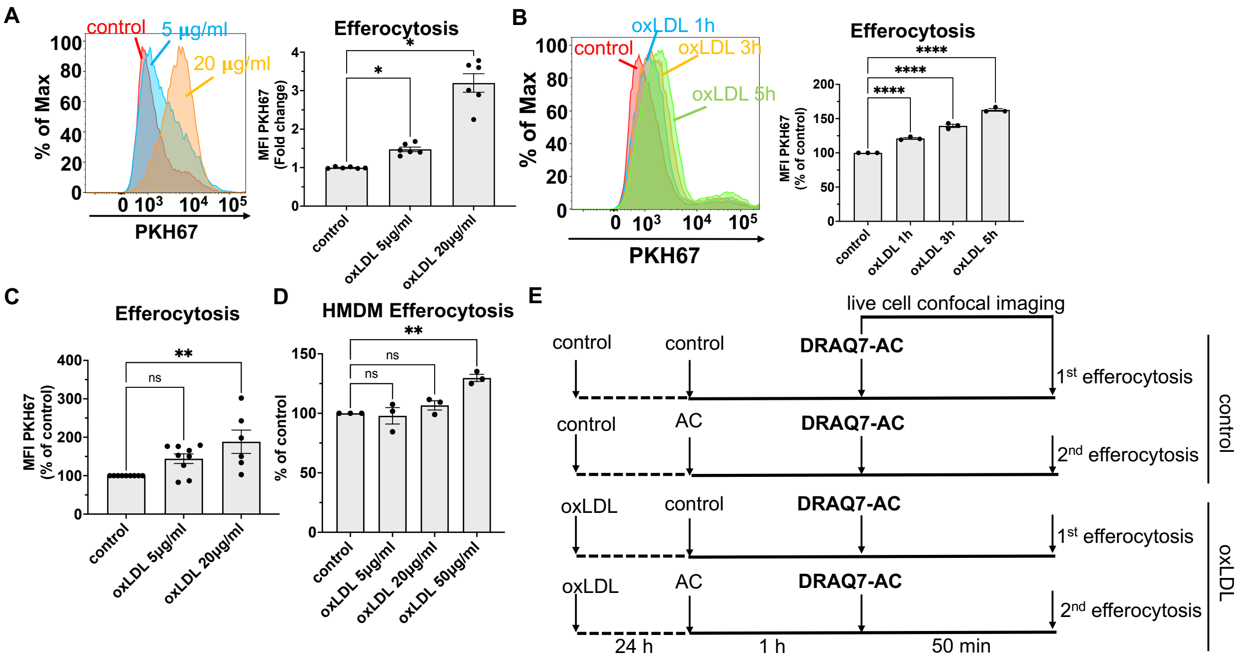


**Figure S2. OxLDL stimulates efferocytosis *in vitro*.** **A**, Examples of histograms of PKH67 fluorescence in WT peritoneal macrophages pre-treated with 5 or 20 µg/ml of oxLDL for 24 h before efferocytosis assay. MFIs are shown in the bar graph; n=6 per group. **B**, WT peritoneal macrophages pre-treated with 20 µg/ml oxLDL for indicated time before efferocytosis assay. MFIs are shown in the bar graph; n=3 per group. **C**, WT peritoneal macrophages pre-treated with 5 or 20 µg/ml of oxLDL for 24h before efferocytosis assay by co-incubation with neutrophil-derived AC. MFIs are shown in the bar graph; n=6-8 per group. **D**, HMDMs were pre-treated with 5, 20 or 50 µg/ml of oxLDL for 24 h before efferocytosis assay. MFIs are shown in the bar graph; n=3 per group. **E**, The schematic experimental design of live cell imaging for the continual efferocytosis assay with or without oxLDL pre-treatment. Max, maximum fluorescence intensity. ns, not significant; *, p<0.05; **, p<0.01; ****, p<0.0001.

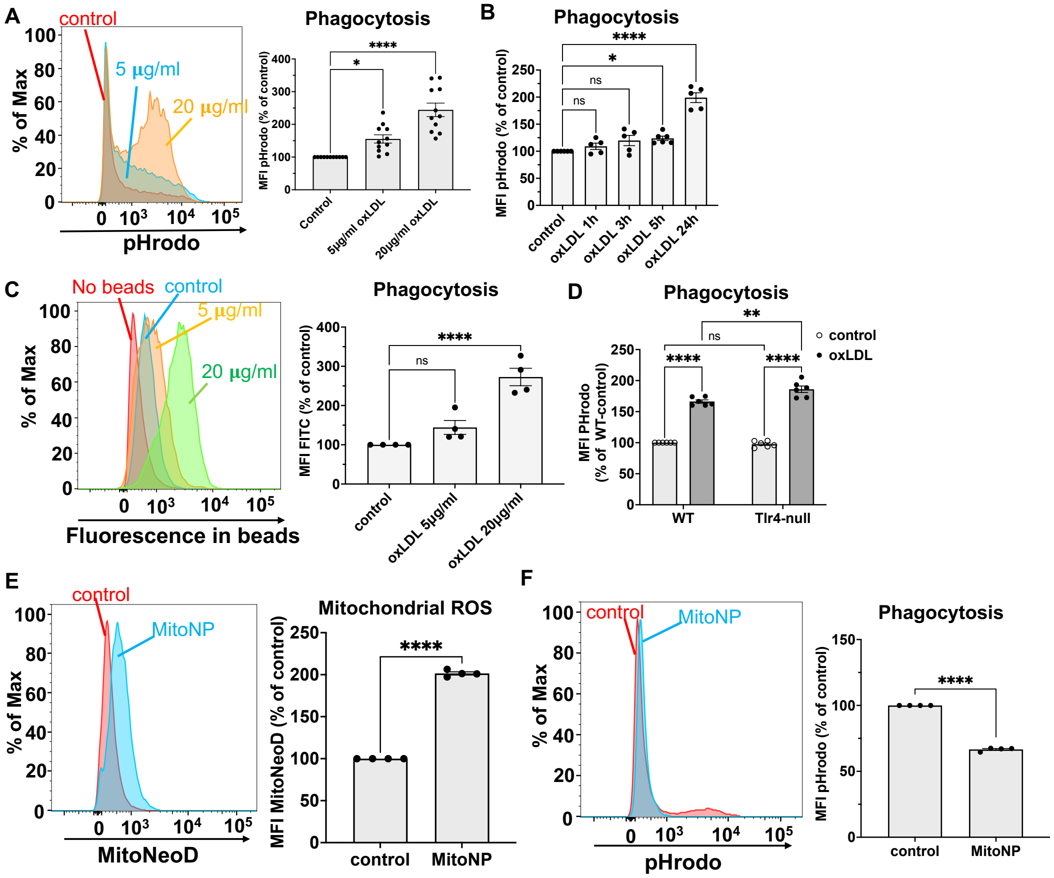


**Figure S3. OxLDL stimulated phagocytosis, but ETC inhibition or mtROS induction is insufficient to stimulate phagocytosis. A**, Examples of histograms of pHrodo fluorescence in WT peritoneal macrophages pre-treated with 5 or 20 µg/ml of oxLDL for 24 h before phagocytosis assay. MFIs are shown in the bar graph; n=4 per group. **B**, WT peritoneal macrophages pre-treated with 20 µg/ml oxLDL for indicated time before phagocytosis assay. MFIs are shown in the bar graph; n=3 per group. **C**, Examples of histograms of bead fluorescence in WT peritoneal macrophages pre-treated with 5 or 20 µg/ml oxLDL for 24h, followed by 10 µm green fluorescence beads co-incubation for 16 h (phagocytosis assay) and flow cytometry analysis. MFIs are shown in the bar graph; n=4 per group. **D**, WT or *tlr4*-null peritoneal macrophages were pre-treated with 20 µg/ml oxLDL for 24 h before co-incubation with pHrodo-conjugated *E. coli* bioparticles (phagocytosis assay). MFIs are shown in the bar graph; n=3 per group. **E**, WT peritoneal macrophages were treated with 10 µM of mitochondrial-targeted redox cycler, MitoNP, for 5 h. Examples of histograms of MitoNeoD fluorescence were shown, and MFIs are shown in the bar graph; n=4 per group. **F**, the same treatment as in E, examples of histograms of pHrodo fluorescence were shown, and MFIs are shown in the bar graph; n=4 per group. ns, not significant; *, p<0.05; **, p<0.01; ****,

$p<0.0001$.

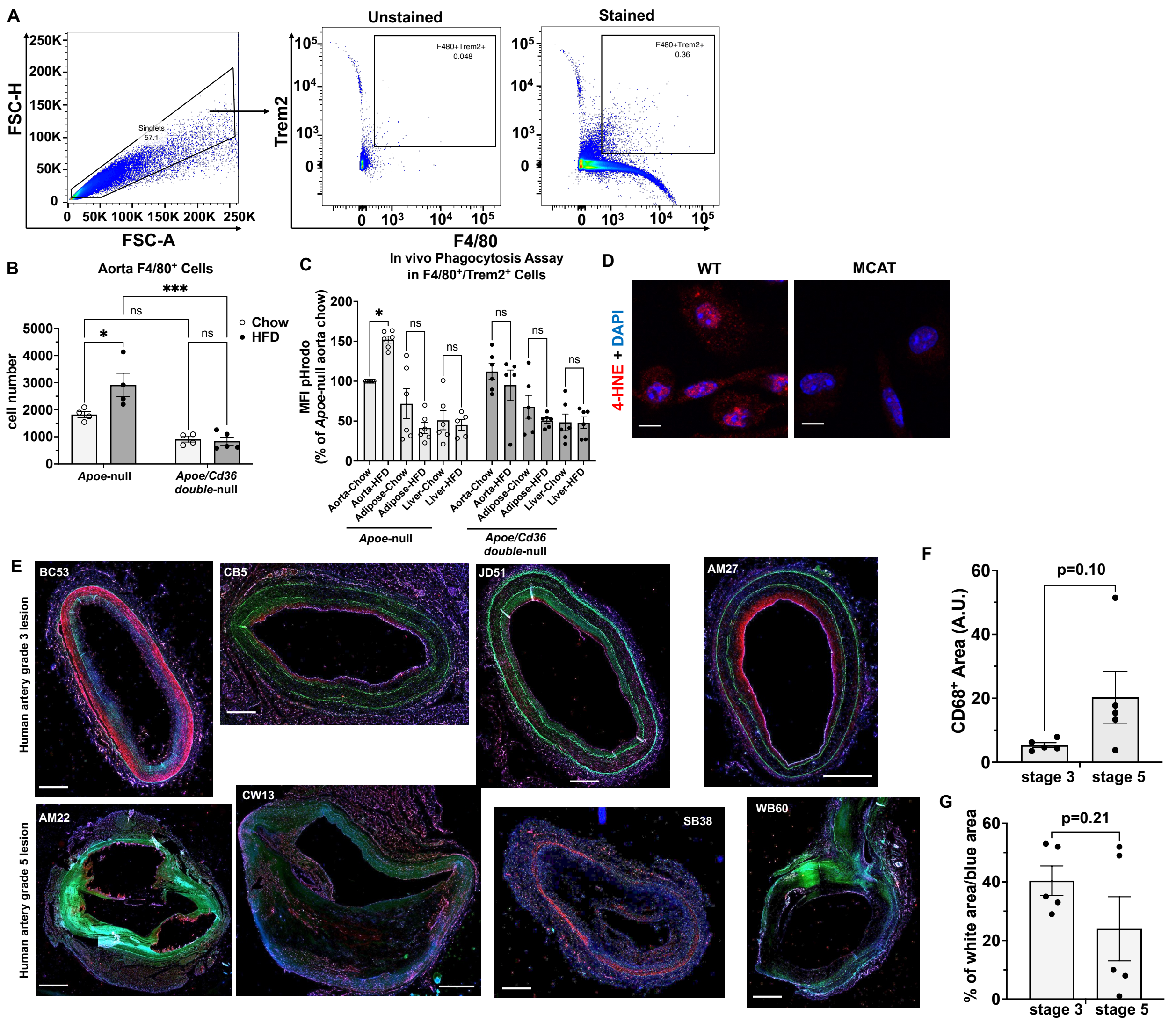

**Figure S4. Association of ROS signaling and phagocytosis/efferocytosis in early stages of atherosclerosis. A**, The gating strategy for selecting F4/80$^+$/Trem2$^+$ macrophages using flow cytometry. Cell clumps were excluded for further analysis using the FSC-A vs. FSC-H strategy. Then F4/80$^+$/Trem2$^+$ macrophages were identified based on a comparison of stained samples vs unstained samples. **B**, Total amount of aortic F4/80$^+$ macrophages were quantified and shown in the bar graph; n=4-5 individual mice per group. **C**, Aortas, adipose tissues, and livers are removed and digested into single-cell suspension for *in vivo* phagocytosis assay. Different tissue cells were then stained by anti-F4/80, anti-Trem2 antibodies. pHrodo MFI was quantified in different tissue-associated F4/80$^+$/Trem2$^+$ macrophages and shown in the bar graph; n=5-6 individual mice per group. **D**, WT or MCAT peritoneal macrophages were treated with 20 µg/ml oxLDL for 24 h and stained with 4-HNE (red). Nuclei were stained with DAPI (blue). Representative images are shown. Scale bar, 10 µm. **E**, Human artery cross sections were co-stained with 4-HNE (red), CD68 (blue), and TUNEL (green). The upper 4 images were from stage 3 lesions, and the bottom 4 images were from stage 5 lesions. Scale bar: 500 μm. **F**, CD68$^+$ (blue) areas from all 10 human lesion samples were quantified and shown in the bar graph. **G**, Three-color colocalized (white) areas, which indicate ROS-associated efferocytes were quantified, normalized by total CD68$^+$ area and shown in the bar graph. ns, not significant; *, p<0.05; ***, p<0.001.

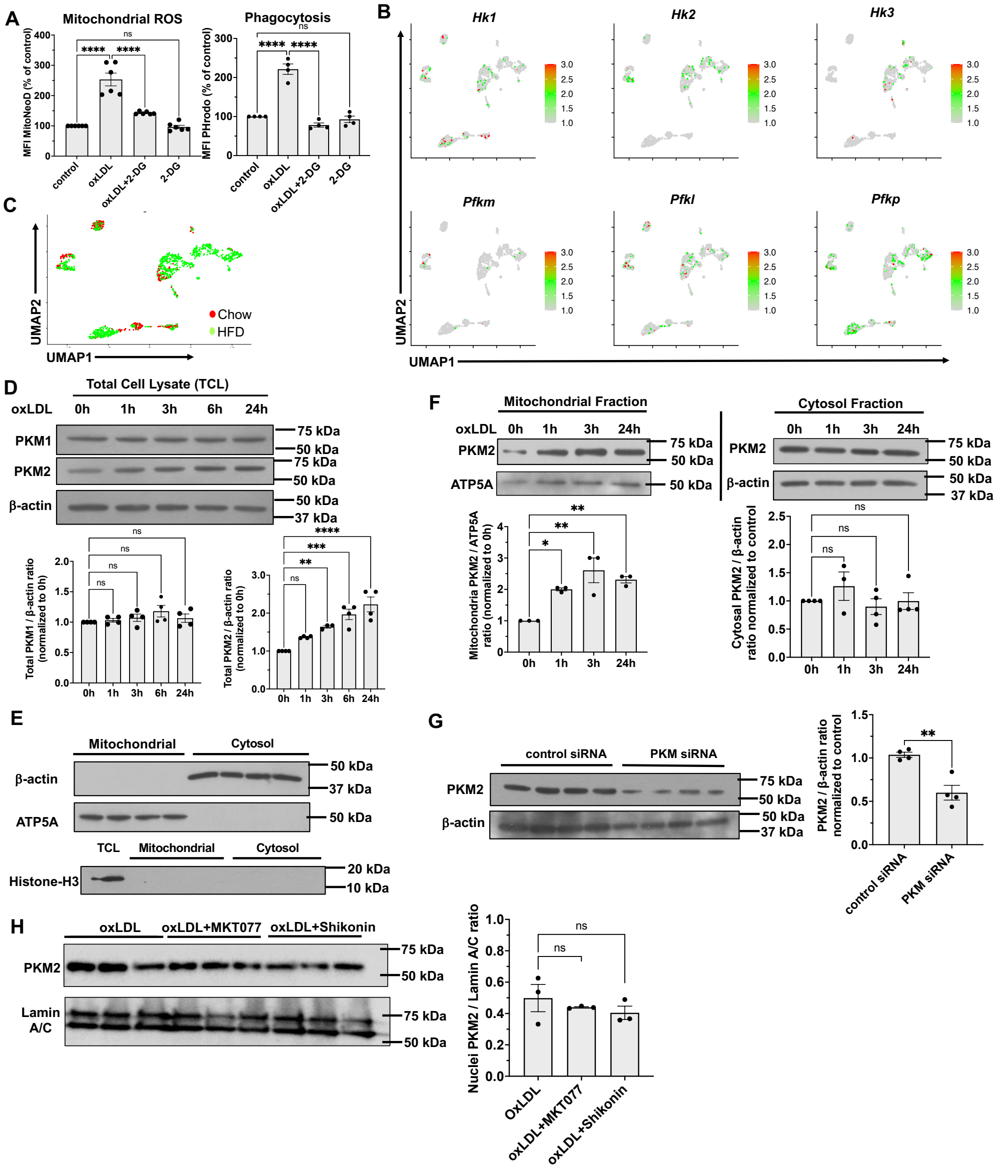


**Figure S5. The glycolytic enzyme PKM2 translocates to the mitochondria. A**, WT peritoneal macrophages pre-treated with 20 μg/ml oxLDL or in combination with 10 mM 2-DG for 24h before mtROS or 3h before phagocytosis assay. The MitoNeoD (left) or pHrodo (right) MFI was quantified and shown in the bar graph; n=3 per group. **B**, Mouse scRNA-seq data were re-analyzed as described for Figure 5. The expression pattern of the genes (Hk1, Hk2, Hk3, Pfkm, Pfkl, Pfkp) encoding two rate-limiting steps of glycolysis is shown in the UMAP. **C**, UMAP representation of cellular origins (chow diet vs HFD 11-week). **D**, WT macrophages were treated with 20 μg/ml oxLDL for indicated time periods. Representative Western blot images of PKM1, PKM2, and β-actin (loading control) were shown. Images were quantified and expressed as fold change of 0 h (control). n=4 per group. **E**, Representative Western blot images of β-actin (cytosol marker), ATP5A (mitochondria marker) and Histone-H3 (nuclei marker) were shown from cell fractions. TCL: total cell lysate. **F**, WT macrophages were treated with 20 μg/ml oxLDL for indicated time periods before cell fractionation. PKM2 and ATP5A (mitochondria fraction loading control) blot images from mitochondrial fractions were shown on the left. PKM2 and β-actin (cytosol fraction loading control) blot images from cytosol fractions were shown on the right. Images were quantified and expressed as fold change of control. n=3-4 per group. **G**, HMDMs were transfected with PKM siRNA for 24 h before lysis. Representative Western blot images of PKM2 and β-actin were shown. Images were quantified and expressed as fold change of control. n=4 per group. **H**, WT macrophages were treated with 20 μg/ml oxLDL or in combination with 0.1 μM MKT077 or 1 μM shikonin for 24 h. Nuclei fractions of the cells were subjected to immunoblot of PKM2 and Lamin A/C (loading control). Images were quantified and shown in the bar graphs. ns, not significant; *, $p<0.05$; **, $p<0.01$; ***, $p<0.001$; ****, $p<0.0001$.

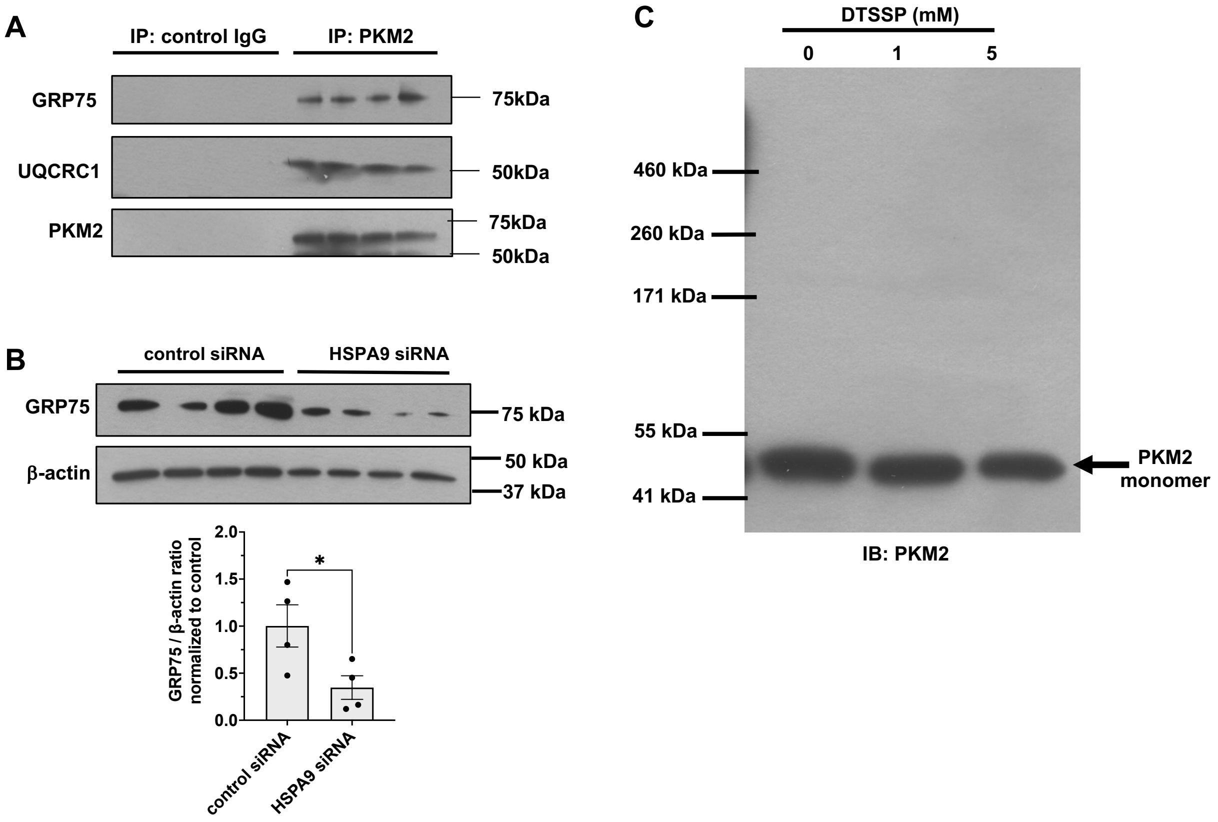


**Figure S6. A,** Examples of Western blot images of GRP75, UQCRC1, and PKM2 from control IgG and anti-PKM2 IgG immunoprecipitates demonstrating specific bindings of PKM2 to GRP75 and UQCRC1. **B**, HMDMs were transfected with HSPA9 siRNA for 24 h before lysis. Representative Western blot images of GRP75 and β-actin were shown. Images were quantified and expressed as fold change of control. n=4 per group. **C**, Purified His-tagged human PKM2 proteins were incubated with DTSSP crosslinker for 1 h at room temperature, followed by SDS-PAGE and immunoblot for PKM2. A representative blot image is shown. n=5.

**SUPPLEMENTAL TABLES**

| Label | Site | Grade | Stable? | Race? | Sex | Age | C.O.D. |
|---|---|---|---|---|---|---|---|
| BC53 | Rco | 3 | | W | F | 53 | Not yet reported |
| CB5 | LAD | 3 | | N.D. | M | 22 | Idiopathic cardiomegaly |
| JD51 | Rco | 3 | | W | M | 48 | Not yet reported |
| JM52 | LAD | 3 | | W | F | 30 | Blunt Force Trauma |
| AM27 | rco | 3 | | W | F | 34 | Cirrhosis |
| AM22 | LAD | 5 | Unstable | W | M | 55 | Bronchal Pneumonia/cirrhosis |
| CW13 | Rco | 5 | Unstable | N.D. | F | 60 | Hypertensive intercranial hemorage |
| JB25 | LAD | 5 | Unstable | W | M | 50 | pulmonary embolus |
| SB38 | rco1 | 5 | Unstable | N.D. | F | 48 | HTN/ASCVD |
| WB60 | RCo | 5 | Unstable | N.D. | N.D. | N.D. | Not determined |

**Table S1.** Basic information on human artery samples. Rco: origin portion of right coronary artery; LAD: left anterior descending artery; N.D.: not determined; C.O.D.: cause of death.

| | Protein name | control intensity | oxLDL intensity |
|---|---|---|---|
| 1 | Slc25a12 | 0.0 | 5.0 |
| 2 | Tufm | 0.0 | 3.0 |
| 3 | Uqcrc1 | 0.0 | 2.0 |
| 4 | Ndufs3 | 0.0 | 1.0 |
| 5 | Acadm | 0.0 | 1.0 |
| 6 | Acaa2 | 0.0 | 1.0 |
| 7 | Cyc1 | 0.0 | 1.0 |
| 8 | Letm1 | 0.0 | 1.0 |
| 9 | Ndufs1 | 1.0 | 7.0 |
| 10 | Slc25a13 | 1.0 | 5.0 |
| 11 | Idh2 | 5.0 | 15.0 |
| 12 | Grp75 | 2.0 | 6.0 |
| 13 | Atp5c1 | 2.0 | 5.0 |
| 14 | Decr1 | 2.0 | 5.0 |
| 15 | Uqcrc2 | 2.0 | 4.0 |
| 16 | Cpt1a | 2.0 | 4.0 |
| 17 | Sqor | 8.0 | 13.0 |
| 18 | Slc25a11 | 2.0 | 3.0 |
| 19 | Slc25a1 | 2.0 | 3.0 |
| 20 | Mdh2 | 19.0 | 23.0 |
| 21 | Slc25a5 | 12.0 | 11.0 |
| 22 | Hadhb | 30.0 | 22.0 |
| 23 | Hars2 | 7.0 | 5.0 |
| 24 | Ndufv1 | 1.0 | 0.0 |
| 25 | Nnt | 1.0 | 0.0 |
| 26 | Cyp11a1 | 1.0 | 0.0 |

**Table S2.** List of 26 mitochondrial proteins co-IP with PKM2. Two (UQCRC1 and GRP75, highlighted in red color) were picked for validation and further analysis.

**Legends for Supplemental Videos**

**Video S1:** This 3D-structured image illustrates the interaction between WT murine peritoneal macrophages (green) and apoptotic cells (ACs, purple) following the primary efferocytosis protocol (described in Figure S2E). Macrophages were cultured in full culture medium (control). At the conclusion of the efferocytosis assay, confocal z-stack images were captured and reconstructed to create this 3D representation.

**Video S2:** This 3D-structured image illustrates the interaction between WT murine peritoneal macrophages (green) and apoptotic cells (ACs, purple) following the secondary efferocytosis protocol (described in Figure S2E). Macrophages were cultured in full culture medium (control). This 3D representation was generated using the same method as Video S1.

**Video S3:** This 3D-structured image illustrates that WT murine peritoneal macrophages (green) engaged with multiple apoptotic cells (ACs, purple) following the primary efferocytosis protocol (described in Figure S2E). Macrophages were pretreated with 20 μg/ml oxLDL for 24 h. This 3D representation was generated using the same method as Video S1.

**Video S4:** This 3D-structured image illustrates the interaction between WT murine peritoneal macrophages (green) and apoptotic cells (ACs, purple) following the secondary efferocytosis protocol (described in Figure S2E). Macrophages were pretreated with 20 μg/ml oxLDL for 24 h. This 3D representation was generated using the same method as Video S1.

**Video S5:** This time-lapse video shows WT murine peritoneal macrophages (pointed by red arrows) interacting with apoptotic cell bodies (ACs, pointed by yellow arrows) in full culture medium (control). Images were captured every 15 seconds for 45 minutes using a NanoLive CX-A label-free Live Cell Imaging System and compiled into a 24-second video. The left macrophage demonstrates sequential uptake of ACs. The right macrophage engages with one AC (top right corner) while ignoring another that appears at 13 seconds into the video. This footage illustrates the normal behavior of macrophages during efferocytosis.

**Video S6:** This time-lapse video shows oxLDL-pretreated (20 μg/ml for 24h) WT murine peritoneal macrophages (pointed by red arrows) interacting with AC (pointed by yellow arrows). Images were captured using the same method as in Video S5 and compiled into a 23-second video. The macrophages exhibited a distorted morphology with thinner and longer filopodia. They appeared to engage multiple ACs simultaneously. These phenotypes differ markedly from those of control macrophages (see Video S5). This footage illustrates that oxLDL alters the macrophage morphology and efferocytosis behavior.
